# Physiological responses to light explain competition and facilitation in a tree diversity experiment

**DOI:** 10.1101/845701

**Authors:** Shan Kothari, Rebecca Montgomery, Jeannine Cavender-Bares

## Abstract

- Ecologists often invoke interspecific facilitation to help explain positive biodiversity-ecosystem function relationships in plant communities, but seldom test how it occurs. One mechanism through which one species may facilitate another is by ameliorating abiotic stress. Physiological experiments show that a chronic excess of light can cause stress that depresses carbon assimilation. If shading by a plant’s neighbors reduces light stress enough, it may facilitate that plant’s growth. If light is instead most often a limiting factor for photosynthesis, shading may have an adverse, competitive effect.
- In a temperate tree diversity experiment, we measured stem growth rates and photosynthetic physiology in broadleaf trees across a gradient of light availability imposed by their neighbors. At the extremes, trees experienced nearly full sun (monoculture), or were shaded by nearby fast-growing conifers (shaded biculture).
- Most species had slower growth rates with larger neighbors, implying a net competitive effect. On the other hand, the two most shade-tolerant species (*Tilia americana* and *Acer negundo*) and the most shade-intolerant one (*Betula papyrifera*) had faster stem growth rates with larger neighbors. The two most shade-tolerant species had large increases in photoinhibition (reduced dark-acclimated F_v_/F_m_) across the gradient of increasing light availability, which suggests they are more vulnerable to chronic light stress. While most species had lower carbon assimilation rates in the shaded biculture treatment, *T. americana* had rates up to 25% higher. *T. americana* also dropped its leaves 3-4 weeks earlier in monocultures, curtailing its growing season. We conclude that although large neighbors can cause light limitation in shade-intolerant species, they can also increase growth through abiotic stress amelioration in shade-tolerant species. Finally, in shade-intolerant *B. papyrifera*, we find a pattern of stem elongation in trees with larger neighbors, which may suggest that a shade avoidance response accounts for the apparent positive trend in stem volume.
- Synthesis: Both positive and negative species interactions in our experiment can be explained in large part by the photosynthetic responses of trees to the light environment created by their neighbors. We show that photosynthetic physiology can help explain the species interactions that underlie biodiversity-ecosystem function relationships. The insights that ecologists gain by searching for such physiological mechanisms may help us forecast species interactions under environmental change.

## Introduction

The sun’s light powers photosynthesis—the chain of reactions through which plants turn carbon dioxide into the living matter they use to grow and reproduce. A common view in plant ecology is that light is often a limiting resource, such that competition for scarce light can govern the fate of plant species (Braun-Blanquet 1932; Canham et al. 1994; Monsi & Saeki 2005; Dybzinski & Tilman 2007; Hautier et al. 2009). But physiologists have also shown that plants often absorb light beyond their capacity to use it for photosynthesis (Long et al. 1994; Demmig-Adams & Adams 1992). Beyond an initial linear rise with increasing light, photosynthesis begins to saturate, and additional light contributes little to the plant’s carbon gain. Any factor that limits photosynthesis can exacerbate the excess of light, including photosynthetic downregulation and environmental stresses like water limitation, cold temperatures, or nutrient-poor conditions (Fig. 1; Long et al. 1994; Demmig-Adams & Adams 1992).

**Fig. 1.**
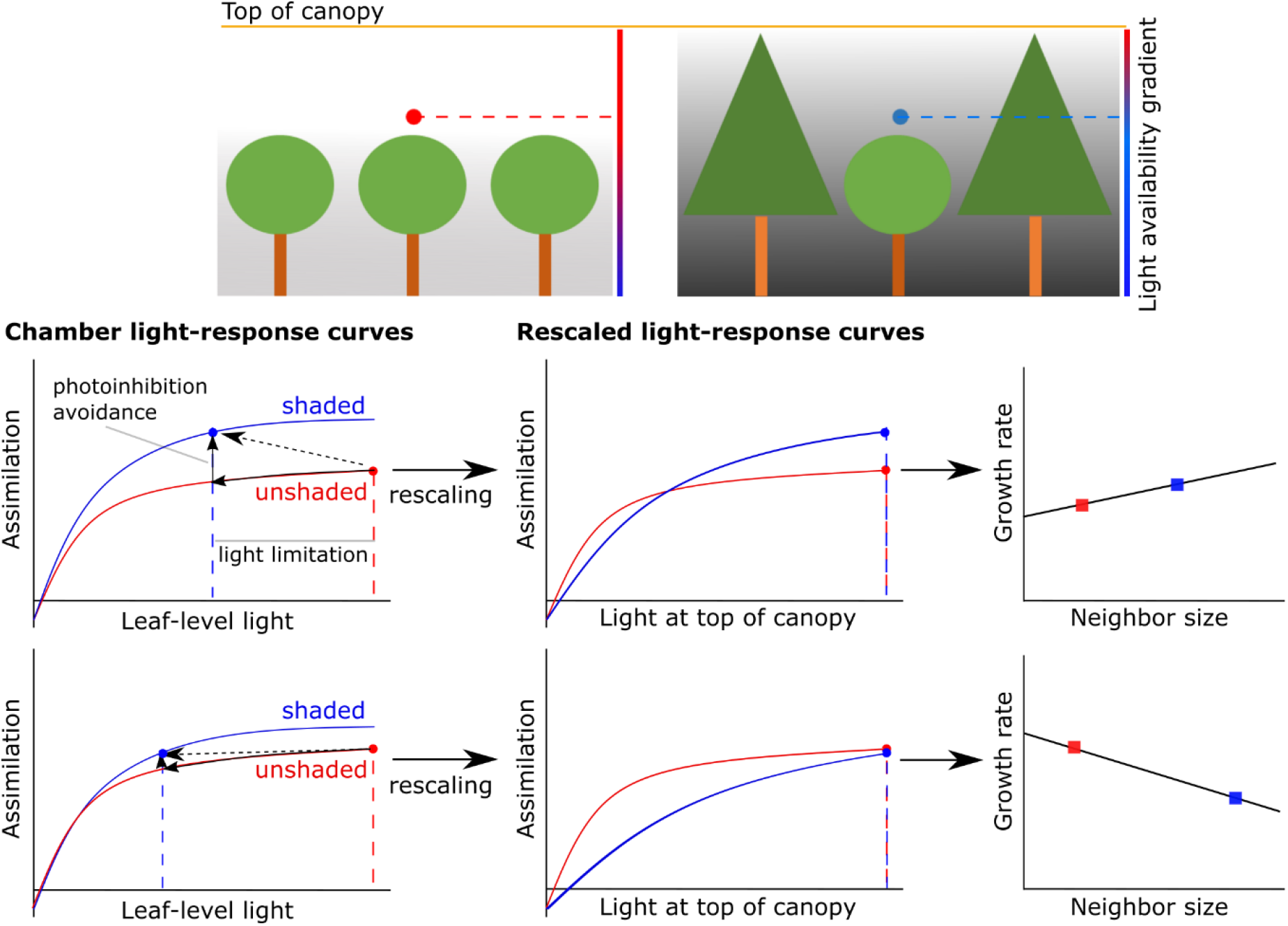
top: Imagine the same slow-growing broadleaf species (round) found in two communities: a monoculture (left), and a biculture (right) with a faster growing conifer (triangular). In the absence of shading by neighbors, all trees would receive the amount of light available at the top of the canopy (yellow line). We define the relative light availability (RLA) for a tree as the fraction of this light that the tree’s top leaves receive. In monoculture, most broadleaf trees have an RLA close to 1. In shaded biculture, they are shaded by their larger conifer neighbors, so their RLA is much less than 1. These trees may be more likely to suffer from light limitation, but they are also at less risk for photoinhibition. Middle: For a single leaf, carbon assimilation tends to increase along a saturating curve as light intensity at the leaf surface increases. Consider two hypothetical photosynthetic light-response curves (left) of leaves from shaded (blue) and unshaded (red) plants. Given a light-response curve, a reduction in light reduces photosynthesis (represented by the small black arrow along the red curve). But if a long-term reduction in light spares the plant from photoinhibition, it can change the shape of the curve such that carbon assimilation at any given light level is greater (represented by the small black vertical arrow). The balance (represented by the large dashed arrow) may either increase or reduce carbon assimilation. In this framework, trees benefit most from shading when light is otherwise abundant: the risk of photoinhibition is greater, and the proportional cost of a given percentage decline in light is lower, since light-response curves are usually concave-down. Whatever the amount of light at the top of the canopy, the top leaves of the shaded plant only receive a corresponding fraction (blue dashed line), while the unshaded plant receives the full amount (red dashed line). To make it easy to compare assimilation rates as a function of light at the top of the canopy, we can rescale the blue light-response curve along the *x*-axis by the inverse of its RLA, making the two dashed lines coincide (middle). In this case, after rescaling, the shaded plant has higher rates of photosynthesis except when the amount of light at the top of the canopy is low. When shaded plants have higher rates of photosynthesis summed over time, they may have faster growth, so growth rates increase with larger neighbors (right). Bottom: We can imagine another tree species (left) in which photoinhibition has a smaller effect (the shaded and unshaded curves are more similar) and the RLA in the shade is lower (the blue dashed line is at a lower light level), such that shading has greater costs and smaller benefits. Here, rescaling makes it clear that the shaded plant has lower photosynthesis at any given level of light at the top of the canopy (middle), so plants shaded by large neighbors may have lower growth rates (right).

Light is thus both an essential resource and a potential stressor. A chronic excess of light can cause plant cells lasting oxidative damage, especially to Photosystem II (PSII). Such damage reduces the efficiency of photosynthesis and can be costly to repair, but plants can avoid it using mechanisms of photoprotection (Murchie & Niyogi 2011). These mechanisms include biochemical pathways that safely dissipate excess light as heat—most notably, the xanthophyll cycle. They also include structural means to avoid absorbing too much light, such as self-shading canopies, reflective leaves, or steep leaf angles (Lovelock & Clough 1992; Streb et al. 1997; Pearcy et al. 2005; Kothari et al. 2018). In general, plants use such photoprotective mechanisms more under environmental conditions that put them at high risk of damage, including high light (Montgomery et al. 2008) and cold or dry climate (Cavender-Bares 2007; Savage et al. 2009; Wujeska et al. 2013; Ramírez-Valiente et al. 2013). Under such conditions, plants need strong photoprotection to prevent even more costly damage.

Depending on how well-protected a plant is, prolonged exposure to excess light can still cause enough damage to reduce carbon assimilation rates (Egerton et al. 2000; Howell et al. 2002; Murchie & Niyogi 2011). Consistent with Long et al. (1994), we use the term ‘photoinhibition’ to describe a drop in carbon assimilation caused by either photodamage or sustained biochemical photoprotection. Bright sunlight causes photoinhibition directly by exposing leaves to more radiation, but it also acts indirectly by altering other microclimatic conditions like temperature, soil moisture, and vapor pressure deficit (VPD; Björkman & Powles 1984). For example, high VPD can cause stomata to close, which limits the use of light for photosynthesis; without an energetic sink, the resulting excess of light can then cause photoinhibition, which limits photosynthetic capacity even further. Alternately, limitation in water or other resources may arrest tissue expansion and cause carbon sink limitation (Tardieu et al. 2014); in response, plants may downregulate photosynthesis and initiate sustained photoprotection (Adams III et al. 2013). Such scenarios may be worsened by belowground competition for resources. Because such stresses vary along environmental gradients, photoinhibition and photoprotective strategies could affect the distribution and persistence of plant populations (Külheim et al. 2002).

Given that excess light can cause stress, it may be that one plant species can facilitate another through shading. Whether a plant receives net costs or benefits from its neighbors’ shade depends on the net balance resulting from (1) the increasing frequency and severity of light limitation and (2) the avoidance of chronic light stress. We can describe these effects in the terms of Monteith et al. (1977), who defined light-use efficiency (LUE) as net photosynthesis divided by the amount of photosynthetically active radiation (PAR) absorbed. Consider a typical light-response curve that plots photosynthesis (*y*-axis) against PAR (*x*-axis; Fig. 1). Chronic photoinhibitory stress may change the shape of the curve by lowering LUE, and thus photosynthesis, at any given amount of PAR. By relieving photoinhibition, shading may raise LUE. But shading would also move the *x*-coordinate leftward along the curve at any given moment, raising the chance the leaf will be in the curve’s light-limited portion. Whether shade helps or hurts depends on whether, integrated across time, the gain of avoiding chronic stress exceeds the loss caused by potential light limitation.

The species interactions that result from shading could help to explain the results of experiments that manipulate biodiversity, including tree diversity. Experimental and observational research alike has shown that biodiversity often promotes ecosystem functions such as productivity (Tilman et al. 2014). In forests, this trend has been shown in both experimental (Potvin & Gotelli 2008; Tobner et al. 2016; Grossman et al. 2017; Williams et al. 2017; Huang et al. 2018; Zemp et al. 2019) and observational (Gamfeldt et al. 2013; Liang et al. 2016; Oehri et al. 2017) data, across temperate (Tobner et al. 2016; Grossman et al. 2017; Williams et al. 2017), subtropical (Huang et al. 2018), and tropical (Potvin & Gotelli 2008; Zemp et al. 2019) biomes. These positive relationships are robust to variation in climate, successional stage, and other factors (reviewed in Grossman et al. 2018; Ammer 2018).

One of the potential drivers of positive biodiversity-productivity relationships is interspecific facilitation (Wright et al. 2017; Barry et al. 2018). In biodiversity experiments, patterns of productivity within species or functional groups can sometimes serve as evidence for facilitation (Mulder et al. 2001; Fichtner et al. 2017), but they cannot tell us the physiological mechanisms that cause one species to facilitate another. Ecologists may need to understand these mechanisms to forecast species interactions and productivity under environmental change (Wright et al. 2017; Barry et al. 2018).

Here, we tested whether the shade created by tree crowns can facilitate the carbon assimilation and growth of neighboring trees in the Forests and Biodiversity (FAB) experiment at Cedar Creek Ecosystem Science Reserve (East Bethel, MN). We asked: Do some species aid others by shielding them from stress caused by intense sunlight? In particular, we sought to test whether slow-growing, shade-tolerant broadleaf trees might be facilitated by adjacent fast-growing conifers (Table 1). We addressed this question by measuring the physiology and woody growth of broadleaf species across plots that vary in species composition. If certain species do benefit from the shade of their neighbors, we would expect individuals with larger neighbors to have higher growth and carbon assimilation rates. We also posed the hypothesis that trees exposed to high light would use structural and biochemical photoprotective mechanisms more than trees shaded by their neighbors. If true, this finding would imply that trees growing in high light must invest in photoprotection to avoid damage, reinforcing the claim that light stress influences the interactions within these communities.

**Table 1:** Characteristics of species in FAB. Except for *Betula papyrifera*, broadleaf species tended to grow slower than needleleaf conifers. Shade tolerance is drawn from Niinemets and Valladares (2006), who evaluated it along a 1-5 scale where higher values indicate an ability to grow under lower light.

| Scientific name | Species code | Common name | Family | Leaf form | Mean RGR in monoculture, 2013-6 (/year) | Shade tolerance | Partner in shaded biculture |
| --- | --- | --- | --- | --- | --- | --- | --- |
| *Acer negundo* | ACNE | Box elder | Sapindaceae | Broadleaf | 0.039 | 3.47 | None |
| *Acer rubrum* | ACRU | Red maple | Sapindaceae | Broadleaf | 0.379 | 3.44 | *Pinus banksiana* |
| *Betula papyrifera* | BEPA | Paper birch | Betulaceae | Broadleaf | 1.272 | 1.54 | *Pinus banksiana* |
| *Juniperus virginiana* | JUVI | Eastern redcedar | Cupressaceae | Needleleaf | 0.975 | 1.28 | NA |
| *Pinus banksiana* | PIBA | Jack pine | Pinaceae | Needleleaf | 2.288 | 1.36 | NA |
| *Pinus resinosa* | PIRE | Red pine | Pinaceae | Needleleaf | 1.619 | 1.89 | NA |
| *Pinus strobus* | PIST | White pine | Pinaceae | Needleleaf | 1.722 | 3.21 | NA |
| *Quercus alba* | QUAL | White oak | Fagaceae | Broadleaf | 0.735 | 2.85 | *Pinus strobus* |
| *Quercus ellipsoidalis* | QUEL | Northern pin oak | Fagaceae | Broadleaf | 0.722 | NA | *Pinus strobus* |
| *Quercus macrocarpa* | QUMA | Bur oak | Fagaceae | Broadleaf | 0.547 | 2.71 | *Betula papyrifera* |
| *Quercus rubra* | QURU | Northern red oak | Fagaceae | Broadleaf | 0.505 | 2.75 | *Pinus strobus* |
| *Tilia americana* | TIAM | American basswood | Tiliaceae | Broadleaf | 0.724 | 3.98 | *Pinus strobus* |

## Methods

### Experimental design

The FAB experiment was planted in 2013 at Cedar Creek Ecosystem Science Reserve in central Minnesota, USA. Cedar Creek has a continental climate with cold winters and warm summers; the mean annual temperature is about 7 °C and the mean annual precipitation is about 660 mm. The site lies on the Anoka Sand Plain, where upland regions tend to have well-drained, sandy soils (Johnston et al. 1996).

FAB comprises 142 4 × 4 m plots, each planted with 64 trees in a 0.5 × 0.5 m grid. The experiment has three blocks, each with 49 plots arranged in a 7 × 7 square. Neighboring plots within a block share a boundary. Each plot is planted with one- or two-year-old seedlings of 1, 2, 5, or 12 species from a pool of 12 species total, including four evergreen conifers with needle-like leaves and eight winter-deciduous angiosperms with broad leaves (Table 1). Except for five-species plots, each distinct species composition is replicated in each block. All species in a plot have nearly equal frequency and are placed at random. Grossman et al. (2017) describes the experimental design in greater detail. The mean relative growth rate in monoculture during the first three years varied dramatically among species—it was highest in conifers and *Betula papyrifera*, and lowest in *Acer negundo* (Table 1).

### Tree growth

Within each species, we sought to determine how an individual’s stem growth rates were influenced by the size of its neighbors. We assumed that neighbors’ size affects the focal individual’s growth rates, but not vice versa. This assumption of exogeneity was warranted because nearly all of the cross-species variation in growth rates is explained by species identity, so the size of a focal individual’s neighbors is explained much more by their identity than by their interactions with the focal individual.

We surveyed tree growth in late fall of each year. For all living trees, we measured diameter and height to the tallest leader. We measured basal diameter (5 cm from the ground) for trees less than 1.37 m tall and diameter at breast height for trees more than 1.37 m. For the first year that a tree crossed the 1.37 m threshold, we measured both diameters. To predict a tree’s basal diameter when it was not measured, we used species-specific linear relationships (*R*^*2*^ = 0.634–0.784) predicting basal diameter additively from height in the same year and the basal diameter in 2016, which was the last year that the basal diameter of all trees was measured. For the conifer *Pinus banksiana*, basal diameter was measured on few enough trees in 2018 that we could not apply this approach; we instead estimated stem volume under the assumption that relative growth rate (RGR) was the same from 2017 to 2018 as from 2016 to 2017. Because we did not consider *Pinus banksiana* growth as a response variable, this approach avoided circularity and allowed us to get reasonable estimates of size in 2018.

In all other cases, we estimated woody volume as *V* = π*r*^2^ℎ, where *h* is height and *r* is the basal radius. This equation assumes each tree’s woody volume can be approximated as a cylinder, an assumption that has been used and justified in other tree diversity studies in the absence of system-specific allometric equations (Tobner et al. 2016; Williams et al. 2017). We then calculated the RGR (yr^−1^ between 2016 and 2018 as (ln *V*_2018_ − ln *V*_2016_)/2 (Hoffman & Poorter 2002). We chose 2016 as a starting point because no replanting of dead trees occurred after that point. Volume loss could cause RGR to be negative (Mahmoud & Grime 1974), which often occurred when stems or branches broke, or occasionally when plants lost all aboveground tissue and resprouted the following year.

For each tree, we calculated the average woody volume of its eight neighbors (in cardinal and intercardinal directions) in 2018 as a proxy for the intensity of aboveground interactions. Neighbors were sometimes missing because of mortality or because the focal tree was on the experiment’s edge. In such cases, we assigned the neighbor a volume of zero. For each broadleaf species, we tested how individual growth changed as average neighbor size increased using a mixed-effects regression model with plot as a random intercept, implemented in *R v. 3.6.1* (R Core Team 2019) using package *lme4 v. 1.1.2* (Bates et al. 2015). We calculated the marginal *R*^*2*^ (*R*_*m*_^*2*^) of the fixed effect using the method of Nakagawa et al. (2017) as implemented in *MuMIn v. 1.43.17* (Bartoń 2020), and *p*-values using Satterthwaite’s degrees of freedom method as implemented in *lmerTest v. 3.1.2* (Kuznetsova et al. 2017), noting that such values should be interpreted conservatively. We also tested whether any species showed non-monotonic responses to neighbor size using the two-lines test (Simonsohn 2018), without random effects.

Some species showed a positive response in stem growth to neighbor size. Besides facilitation, such a response could also be caused by shade avoidance, which may cause plants to increase shoot biomass at the expense of root biomass in order to compete for light (Shipley & Meziane 2002). We aimed to discern whether such a shade avoidance syndrome could contribute to the positive stem growth response. Lacking root biomass sorted by species, we could not test this idea directly, but we could test another symptom of a typical shade avoidance syndrome: stem elongation, which prioritizes vertical growth over lateral growth (Henry & Aarssen 1999). We considered height and basal diameter in 2016, the last year in which basal diameter was measured directly on all trees. For all species that showed a positive stem growth response, these two variables were linearly correlated. We used mixed-effects linear models with plot as a random intercept to predict diameter from height in each species, then tested whether the residuals—the observed diameters’ deviations from the predictions for a tree of equal height—were associated with average neighbor size. If trees with larger neighbors show a shade avoidance syndrome, we would expect them to have smaller diameters for a given height. We obtained similar conclusions when considering the log-transformed diameter-to-height ratio as the response variable rather than the residuals of the linear diameter-height relationship.

### Photosynthetic physiology

We measured chlorophyll fluorescence parameters and leaf reflectance spectra among all eight broadleaf species, and photosynthetic light-response curves among four focal species (*A. rubrum*, *B. papyrifera*, *Q. ellipsoidalis*, and *T. americana*). For physiological measurements, we focused on plots belonging to one of three treatments: (1) relatively open monocultures of broadleaf species (‘monoculture’); (2) bicultures comprising one broadleaf and one conifer species (‘shaded biculture’); and (3) twelve-species plots (see Fig. S1 for sample images). For each species in each treatment, we measured physiological parameters on six individuals—two in each of three plots. Neither *A. negundo* nor *Q. macrocarpa* was planted with a conifer in any biculture. We used *B. papyrifera* as a shaded biculture partner for *Q. macrocarpa* because it is a fast-growing species and creates shade. We omitted the shaded biculture treatment for *A. negundo*. When possible, we chose the same trees within a plot for each kind of physiological measurement so that we could compare these aspects of physiology within individuals.

#### Photosynthetic light-response curves

We measured photosynthetic light-response curves from the four focal species during July 2018 using an LI-6400 gas exchange system (LI-COR BioSciences, Lincoln, NE, USA) with a 6400-40 Leaf Chamber Fluorometer head. From each tree (*n* = 72 total), we selected a fully expanded upper leaf with no visible sign of disease or herbivory. We noted the angle of each leaf relative to horizontal (to the nearest 15°) before beginning measurements. Each curve had nine steps in descending order of brightness: 2000, 1500, 1000, 500, 200, 100, 50, 20, and 0 μmol m^−2^ s ^−1^. We also measured the relative electron transport rate (ETR) using chlorophyll fluorescence at each light level. We maintained favorable conditions inside the chamber during each measurement. Following each curve, we removed the leaf to measure leaf mass per area (LMA) using a balance and a flatbed scanner. Further details about light-response curve procedures and ETR calculations can be found in the Supplemental Materials (Appendix 1). Finally, we estimated parameters like the light-saturated carbon assimilation rate (*A*_*sat*_) by fitting a non-rectangular hyperbolic model (Johnson & Thornley 1984) to each light-response curve using *R* code written by Nick Tomeo, available at: https://github.com/Tomeopaste/AQ_curves.

We aimed to determine how realized carbon assimilation rates vary with the photosynthetic photon flux density (PPFD) at the top of the canopy, a measure of light availability in the absence of shading. While all trees received the same range of light intensities inside the instrument chamber, some trees received much less light than others *in situ* because their neighbors shaded them more. At any moment, trees that are more shaded are farther to the left on their light-response curve than trees that are less. For example, consider a tree crown whose topmost leaves only receive 50% of the light reaching the top of the canopy. For this tree crown, the realized assimilation rate we estimate at a top of the canopy PPFD of 2000 μmol photons m^−2^ s ^−1^ would be the assimilation rate at a leaf-level PPFD of 1000 μmol m^−2^ s ^−1^. To visualize the impacts of shading by neighbors, we followed the approach of Howell et al. (2002): we estimated how much light each leaf receives as a constant fraction of light at the top of the canopy, and rescaled each leaf’s fitted light-response curve along the *x*-axis by this fraction’s reciprocal. We estimated the constant rescaling factor for each individual as *RLA* × cos*(θ)*, where *RLA* is relative light availability (see *Light availability*) and *θ* is the leaf angle from horizontal. (Taking the cosine of *θ* approximately corrects for the reduction in horizontally projected leaf area in steeply inclined leaves when the sun is directly overhead, as simplified from Ehleringer & Werk [1986].) We used this procedure on each fitted curve in order to compare across treatments in a way that accounts for the varying fractions of light that individuals receive due to differences in relative height and vertical light transmission.

Finally, we used the rescaled light-response curves to estimate carbon assimilation by top leaves throughout July. We used an hourly averaged time series of solar radiation from Carlos Avery Wildlife Management Area in Columbus, MN, 13.6 km away from the study site. Following Udo and Aro (1999), we assumed a conversion rate of 2.08 μmol photons m^−2^ s^−1^ of PAR per Watt m^−2^ of solar radiation. By using these data as inputs to the rescaled light-response curves, we estimated carbon assimilation over the month of July. This procedure assumes that the photosynthetic response to light remains constant and unaffected by factors like leaf temperature and stomatal closure. While this assumption is never quite true, we consider it a useful way of estimating the consequences of realistic fluctuations in light.

We mainly express assimilation per unit of leaf dry mass because we aim to determine the plants’ return on the carbon invested in leaf construction. As expected, leaves growing in low light tended to have lower LMA in most species (Fig. S2; Poorter et al. 2019; Williams et al. 2020). More carbon is spent on constructing a leaf with high LMA, which means that the leaf has to assimilate more carbon to recoup its construction cost. Expressing data on a mass basis accounts for these large differences in construction cost, although we also discuss area-based assimilation rates when they provide additional physiological insight. We also express stomatal conductance (*g*_*s*_) and electron transport rate (ETR) on a mass basis to compare them with mass-based assimilation rates.

#### Instantaneous chlorophyll fluorescence and spectral reflectance

In all eight broadleaf species, we measured chlorophyll fluorescence parameters using an FMS2 pulse-modulated fluorometer (Hansatech Instruments Ltd., Norfolk, UK) over two days in late July (*n* = 138 total). We measured dark- and light-acclimated parameters at the same spot on the same leaves. We attached opaque clips to leaves in the evening before measuring dark-acclimated F_v_/F_m_ within two hours after sunrise. This parameter describes the maximum quantum yield of PSII and is a general index of photoinhibition (Murchie & Lawson 2013). We then took light-acclimated measurements between 12:00 and 14:00 each day. The protocol involved the following steps: actinic light at 1000 μmol m^−2^ s^−1^ for 15 seconds, a saturating pulse, and two seconds of far-red light. We exposed all leaves to the same actinic light because we aimed to assess photoprotective capacity under comparable light conditions, even though each tree had a different light environment.

From these data, we estimated qN, a parameter that indicates how much a plant relies on photoprotective dissipation (non-photochemical quenching) under the imposed actinic light. This parameter is correlated with the de-epoxidation state of xanthophyll cycle pigments (Watling et al. 1997; Cavender-Bares & Bazzaz 2004). Following Kramer et al. (2004), we also calculated the quantum yields ϕ_PSII_, ϕ_NPQ_, and ϕ_NO_, which sum to 1. These three parameters are the proportions of light energy dissipated by photosynthetic chemistry, non-photochemical quenching, and non-regulated dissipation. The last is a crucial parameter because it represents light that a plant cannot dissipate safely; this portion of the absorbed light may contribute to photodamage. The Supplemental Materials contain formulas for all parameters and justifications for our choices in data analysis (Appendix 1).

On separate top leaves of the same trees, we measured reflectance spectra (350-2500 nm) using a PSR+ 3500 field spectroradiometer (Spectral Evolution, Lawrence, MA, USA). We used these spectra to calculate the photochemical reflectance index (PRI), calculated as PRI = (R_531_ - R_570_) / (R_531_ + R_570_), where R_*n*_ is the reflectance at a wavelength of *n* nm. PRI shows a negative correlation with carotenoid: chlorophyll ratios and, over shorter time scales, also tracks the xanthophyll de-epoxidation state (Gamon et al. 1992; Wong & Gamon 2015; Gitelson et al. 2017). We estimated species-specific linear responses of PRI and chlorophyll fluorescence parameters to relative light availability (see *Light availability*).

To understand how photoinhibition and photoprotective strategies vary with adaptations to shade, we used shade tolerance values that Niinemets & Valladares (2006) compiled on a five-point scale. Higher values along this scale indicate that plants are able to grow in lower light conditions.

### Phenology

We monitored the timing of leaf abscission for *T. americana* in monoculture and shaded biculture plots to assess whether shade could delay senescence and abscission, extending the period for carbon gain (Cavender-Bares et al. 2000). In early August, we surveyed 120 *T. americana* trees (60 in monoculture, 60 in shaded biculture) to determine the proportion of leaves that had fallen based on how many of the top five axillary buds lacked a leaf. From each of these trees, we marked a single leaf and returned every 1-2 weeks to monitor its senescence. We assessed each leaf on a binary basis (senesced or not) based on whether it had at least 50% remaining green leaf area. We chose this criterion as a simple proxy for the continuous scale used by Cavender-Bares et al. (2000). Based on the initial survey and the repeated surveys of marked leaves, we tracked the proportion of leaves that had senesced over time. To check whether leaves were photosynthetically active until near the point of senescence, we collected a one-time measurement of dark-acclimated F_v_/F_m_ in mid-September among remaining leaves.

We also performed a one-time measurement of 60 (30 per treatment) *A. rubrum* plants on October 1, using the same protocol we used to do our initial early August survey of *T. americana*. We aimed for this survey to help test whether our results hold across species.

### Water potential

Water deficits—even moderate ones—can arrest tissue growth (Tardieu et al. 2014) and reduce carbon gain by causing stomata to close (Brodribb et al. 2003). To see how our treatments affected water status, we measured leaf water potential using a Scholander pressure bomb. We measured pre-dawn water potential (Ψ_PD_) in all eight broadleaf species (*n* = 96) and midday water potential (Ψ_MD_; between 12:00 and 14:00) in only the four focal species (*n* = 48). We included monoculture and shaded biculture treatments, except in *A. negundo*, where we used the twelve-species treatment in place of the absent shaded biculture treatment. We removed leaves and immediately placed them in impermeable plastic bags with wet paper towels. We carried leaves in coolers to be measured indoors within 90 minutes.

Because we only intended to test whether there was a general tendency across species for water potential to vary with light, we included species as a random intercept and light availability as a fixed effect in mixed-effect regression models for both Ψ_PD_ and Ψ_MD_. We calculated *R*_*m*_^*2*^ and *p*-values as in models for growth rate.

### Leaf angles

Leaf angle is a key control on light exposure because more vertical leaves intercept less light when the sun is directly overhead, and may allow light to penetrate deeper into the canopy (Posada et al. 2009). As a result, steep leaf angles can be a structural strategy to avoid light, temperature, and water stress (Valladares & Pugnaire 1999; Werner et al. 1999; Pastenes et al. 2004; Valiente-Banuet et al. 2010). We predicted that trees in monoculture would have steeper leaf angles than those in other treatments. From each of three species (*A. rubrum*, *Q. ellipsoidalis*, and *T. americana*), we randomly selected five individuals from each of six plots—three from monoculture treatments, three from shaded bicultures (*n* = 15 per species × treatment). On these individuals, we measured the leaf angle from horizontal of the top five leaves in the crown to the nearest 15°.

Because angles of leaves within individuals may be non-independent, we tested whether treatments differ using a mixed-effects model with treatment as a fixed effect and individual tree as a random intercept. Deviations from horizontal in either direction can reduce light interception, so we used the absolute value of leaf angle as the dependent variable. We again calculated *p*-values using Satterthwaite’s degrees of freedom method.

### Environmental data

#### Light availability

On a cloudy day with diffuse light conditions, we measured the available light above each tree selected for this study (*n* = 138) using an AccuPAR LP-80 ceptometer (METER Group, Pullman, WA, USA). We took the mean of two to four PAR measurements in the open and the mean of two measurements directly above the topmost leaf of each tree. By calculating a ratio of these values, we could calculate the percent of light transmitted to the top of each tree. We called this value *relative light availability* (RLA). This value usually correlates well with the percentage of light a tree can access over much longer time-scales (Parent & Messier 1996).

#### Soil water content

We collected a one-time measurement of soil volumetric water content in mid-August using a FieldScout Time-Domain Reflectometer 350 (Spectrum Technologies, Inc., Aurora, IL, USA). This single measurement followed six days from most recent rain. At midday, we collected and averaged three measurements in each of 32 plots, spanning monocultures of the four core broadleaf species, shaded bicultures, and twelve-species plots.

#### Air temperature

For nine days during early September, we set out four Thermochron iButtons (model DS1921G; Maxim Integrated Products, Sunnyvale, CA, USA) in shaded plots and two in twelve-species plots, setting them to log air temperature hourly. Each iButton was suspended on mesh inside a 1 m-tall PVC pipe that was capped with an elbow to shield against solar irradiance. We compared these data to air temperature measured hourly under open, sunny conditions at the Cedar Creek LTER weather station about 0.77 km away. We assumed that data from this station would be representative of air temperature in non-shaded areas. We ignored nighttime and early morning (20:00-08:00 h) readings because they seemed to be influenced by water condensation and evaporation.

## Results

### Tree growth

Species varied in their relationship between neighbor stem volume and focal individual relative growth rate (RGR; mixed-effects ANCOVA, neighbor volume × species interaction; *p* < 10^−15^; F_11,7279_ = 12.440). For individual species, relationships between RGR and neighbor volume were noisy but often highly significant (Fig. 2). In most species, individuals with larger neighbors had lower RGR, including in *A. rubrum* (*R*_*m*_^*2*^ = 0.027, *p* < 0.005, *t*(366) = −3.257), *Q. alba* (*R*_*m*_^*2*^ = 0.030, *p* < 10^−4^, *t*(486) = −3.932), *Q. ellipsoidalis* (*R*_*m*_^*2*^ = 0.051, *p* < 10^−7^, *t*(341) = −5.485), *Q. macrocarpa* (*R*_*m*_^*2*^ = 0.006, *p* = 0.0380, *t*(697) = −2.079), and *Q. rubra* (*R*_*m*_^*2*^ = 0.053, *p* < 10^−7^, *t*(444) = −5.785). But *B. papyrifera* had a weak positive response to neighbor size (*R*_*m*_^*2*^ = 0.010, *p* = 0.049, *t*(124) = 1.993), and two more species grew faster with larger neighbors, but only when average neighbor size was log-transformed: *A. negundo* (*R*_*m*_^*2*^ = 0.014, *p* = 0.019, *t*(380) = 2.361) and *T. americana* (*R*_*m*_^*2*^ = 0.018, *p* < 0.005, *t*(350) = 2.154). These results are qualitatively unchanged by removing trees on a block edge or including focal plant volume in 2016 as a covariate, although the effect of neighbor size in *B. papyrifera* or *Q. macrocarpa* sometimes became statistically insignificant under these alternate model specifications (Supplemental Materials, Appendix 2). No species had a non-monotonic response to neighbor size.

**Fig. 2:**
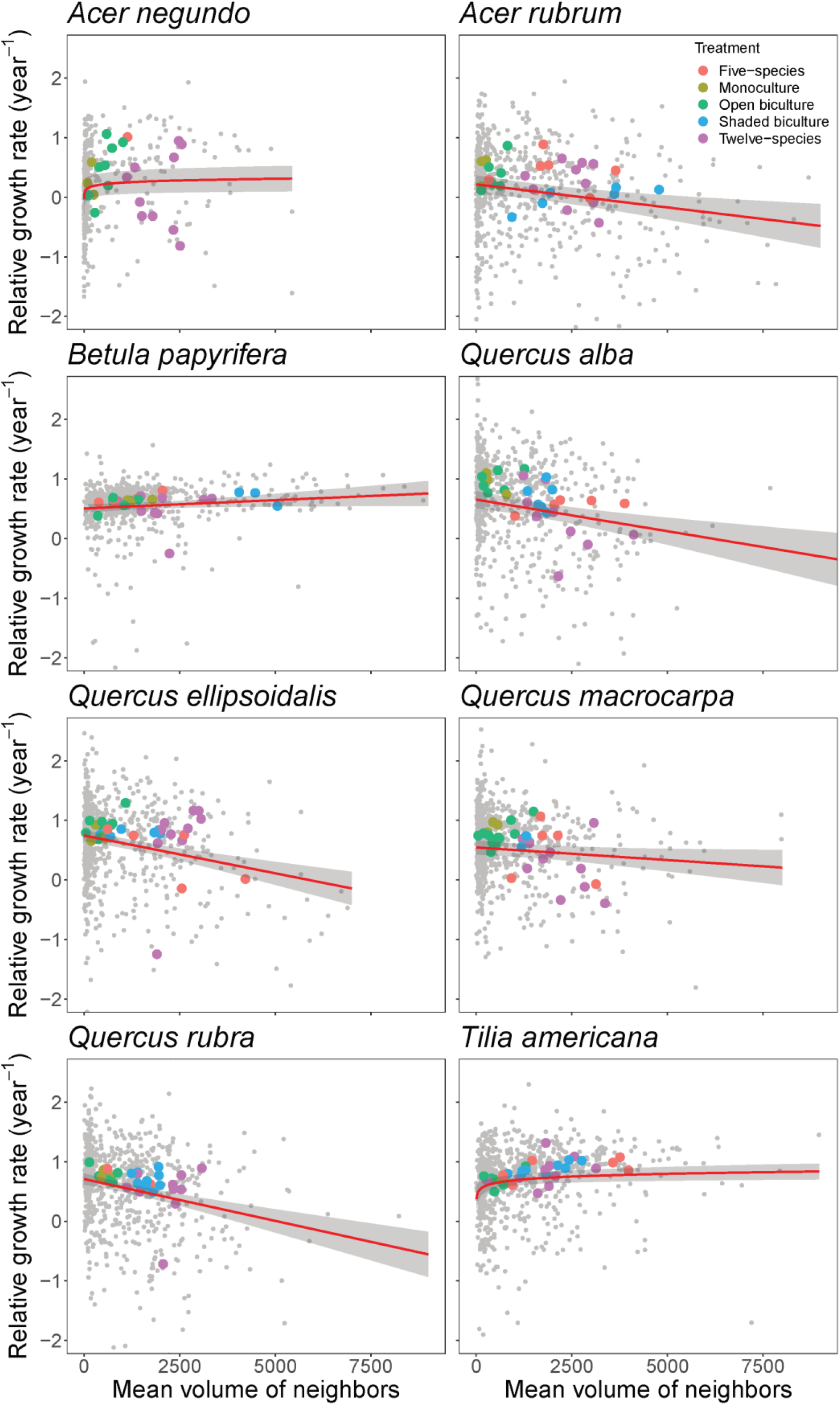
Relative growth rate (RGR) of woody stem volume in individuals of each species between fall 2016 and 2018 as a function of the average stem volume of all neighbors. Gray dots represent individuals, and the regression is fit based on a mixed-effects model with plot as a random intercept. For *Tilia americana* and *Acer negundo*, the relationship with neighbor volume is modeled as logarithmic rather than linear. The large colored dots aggregate data to the plot scale, as described in the main text, and are color-coded by treatment. A shaded biculture is a plot where the focal broadleaf species grows with either a conifer or *Betula papyrifera* (unless *B. papyrifera* is the focal broadleaf species). An open biculture is a plot where the focal broadleaf grows with a broadleaf species other than *B. papyrifera*. About 1.1% of trees fall outside the plot bounds.

Trees that died between 2016 and 2018 dropped out of our individual-level analyses of RGR because their stem volume was 0 in 2018, making it impossible to calculate RGR. To see whether mortality could alter our conclusions, we also performed statistical analyses on absolute change in stem volume from 2017 to 2018, in which we alternately (1) treated mortality as a decline in volume to 0, or (2) removed trees that died from analyses. The rate of mortality was low between 2016 and 2018 (~7.5%), and accounting for mortality makes little qualitative difference when considering absolute growth rate as the response variable (Fig. S3; Supplemental Materials, Appendix 2).

Although the individual-level effects are noisy, they reveal strong tendencies in average stem growth across the full range of neighbor size. Imagine two *T. americana* trees that begin at the same size: One in monoculture, where the average neighbor in 2018 is 370 cm^3^, and one growing in a shaded biculture with *Pinus strobus*, where the average neighbor is 2120 cm^3^. The individual-level regression (with log-transformation) predicts that after two years, the average *T. americana* in the shaded biculture would be 26.7% larger because of the difference in mean RGR. Or consider *Q. rubra*, which had the strongest negative response to neighbor size: We predict that an average tree in the shaded biculture (mean neighbor size 1988 cm^3^) would be 40.2% smaller than in monoculture (mean neighbor size 160 cm^3^). These figures assume that individuals across treatments start at the same size at the initial point for RGR estimates, so they may underrepresent the true differences in size that have developed. The absolute growth rate gives some insight into these differences. For example, the median *T. americana* stem in monoculture grew 99 cm^3^ (25–75^th^ percentile: 21–237 cm^3^) between 2017 and 2018, compared to 267 cm^3^ (29–633 cm^3^) in shaded biculture with *P. strobus* or *J. virginiana*. In comparison, the median *Q. rubra* stem grew 32 cm^3^ (25–75^th^ percentile: 3.9–124 cm^3^) in monoculture, compared to only 12 cm^3^ (−5.2–89 cm^3^) in shaded biculture with *P. strobus*. In general, trends in the absolute growth rate across the gradient of neighbor size mirrored those in the relative growth rate (Supplemental Materials, Appendix 2).

We can also aggregate these observations to the plot scale—for example, by considering the RGR of summed stem volume of each species in each plot from 2016 to 2018 (Fig. 2). (This approach is distinct from an average of individual RGRs, which assigns equal weight to small and large individuals.) Although aggregation leaves less statistical power to detect relationships, it also reduces noise, such that the relationships we find explain much more of the variation in growth across plots. As average neighbor size increases, RGR declines in *Q. alba* (*R*^*2*^ = 0.297, *p* < 0.005, *t*(28) = −3.642) and *Q. macrocarpa* (*R*^*2*^ = 0.265, *p* < 0.005, *t*(31) = −3.540), but increases in *T. americana* (*R*^*2*^ = 0.312, *p* < 0.001, *t*(32) = 3.995).

Finally, for the three species that showed positive individual-level stem growth responses to neighbor size (*T. americana*, *A. negundo*, and *B. papyrifera*), we performed a follow-up analysis to test whether they showed a shade avoidance response. We found that trees with larger neighbors had smaller diameters than expected based on their height in *B. papyrifera* (*R*^*2*^ = 0.009, *p* = 0.008, *t*(795) = −2.645), but not *T. americana* and *A. negundo* (*p* > 0.05).

### Photosynthetic physiology

#### Photosynthetic light-response curves

As a function of the chamber light level, mass-based assimilation rates were always higher in shaded biculture and twelve-species treatments than in monoculture in three out of the four focal species (*Q. ellipsoidalis*, *A. rubrum*, and *T. americana*; Fig. 3, left). In the early-successional species *B. papyrifera*, mass-based assimilation rates in shaded biculture were lower than in the other two treatments. Area-based assimilation rates showed similar trends in *T. americana*, but varied less among treatments in *B. papyrifera* and *Q. ellipsoidalis*—and in *A. rubrum*, they were lower in shaded biculture than the other treatments (Fig. S4) as a result of large differences among treatments in LMA (Fig. S2).

**Fig. 3:**
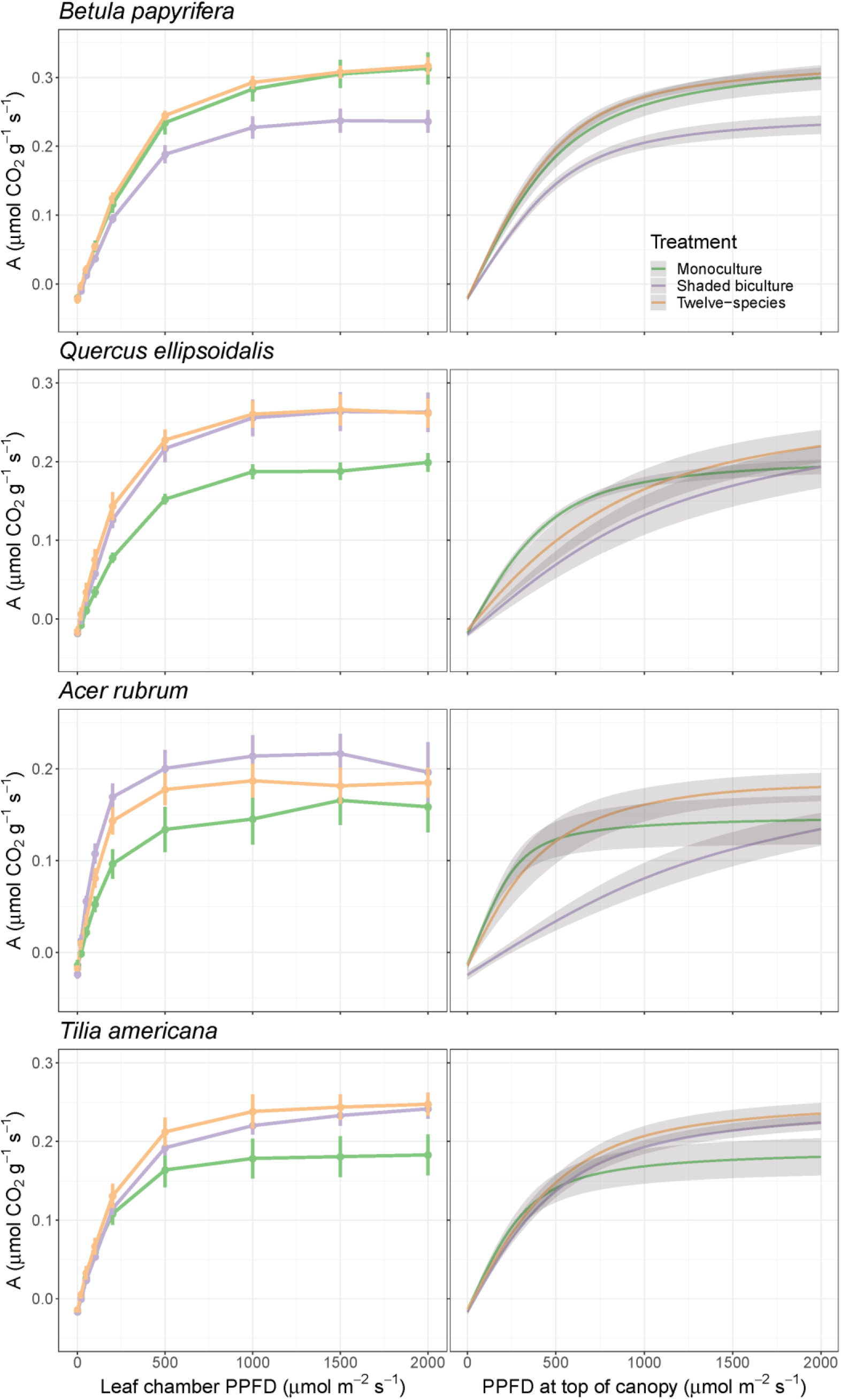
Mass-based photosynthetic light-response curves for four broadleaf species. The left panels depict carbon assimilation rates as a function of chamber light intensity; the right panels present estimates of realized carbon assimilation rates as a function of light intensity at the top of the canopy. These rates are lower because of shading by neighbors (as in Fig. 1, bottom right). Species are arranged from the least shade-tolerant (*Betula papyrifera*) to the most (*Tilia americana*). Error bars in left panels and gray ribbons in right panels are ± 1 SE.

In all broadleaf species except *B. papyrifera* and *T. americana*, LMA was higher in the monoculture than the other two treatments (Fig. S2). Across the four focal species, the light-saturated assimilation rate *A*_*sat*_ on an area-basis was positively correlated with LMA (*R*^*2*^ = 0.431; *p* < 10^−9^; *t*(69) = 7.349). Within species, there was no correlation in *B. papyrifera* and *A. rubrum*, but a positive correlation in *Q. ellipsoidalis* (*R*^*2*^ = 0.192; *p* = 0.039; *t*(16) = 2.243) and *T. americana* (*R*^*2*^ = 0.436; *p* = 0.002; *t*(16) = 3.759). In *T. americana*, the intercept was statistically indistinguishable from zero, making the relationship between area-based *A*_*sat*_ and LMA close to a direct proportionality.

The rate of photosynthesis is often limited either by ETR or by the RuBP carboxylation rate; the latter may in turn be limited by stomatal diffusion of CO_2_ (Farquhar et al. 1980). In our data, both ETR and stomatal conductance (*g*_*s*_) increased with light availability (Fig. S5). (ETR tended to decline at very high light levels, especially in shaded treatments, perhaps because of acute photoinhibition.) Within each species, the rank-order of treatments in carbon assimilation rates, ETR, and *g*_*s*_ were broadly congruent. One exception is that *T. americana* in shaded biculture had much higher *g*_*s*_ than in twelve-species or monoculture plots, despite having slightly lower assimilation rates than in twelve-species plots.

Compared to the unscaled chamber light-response curves, the picture that emerged from rescaled light-response curves is more complex (Fig. 3, right). In *B. papyrifera*, the assimilation rate was still lowest across light levels in the shaded biculture treatment. In late-successional *T. americana*, the assimilation rate was higher in shaded biculture and twelve-species treatments by up to 25% compared to monoculture across most light levels. In the other two species, the mass-based assimilation rate was highest in monoculture at low light, but twelve-species and (in *Q. ellipsoidalis*) shaded biculture plots intersected and surpassed monocultures when enough light was available. *A. rubrum* and *Q. ellipsoidalis* grew less and had lower light availability in shaded bicultures than *T. americana* (Fig. S2), so their mass-based assimilation rates dropped considerably more in the rescaled light-response curves. The area-based assimilation rate in these two species was lower in shaded bicultures (and in *Q. ellipsoidalis*, twelve-species plots) than in monocultures across the full domain of light availability (Fig. S4), but otherwise, mass-based and area-based rates showed similar patterns across treatments within species.

Using a time series of solar radiation, we estimated total mass-based assimilation rates in July 2018 based on each rescaled light-response curve (Fig. S6). We found that trees would have lower total assimilation in shaded biculture than in monoculture in *A. rubrum* (ANOVA; *p* = 0.009; F_2,14_ = 6.759) and *B. papyrifera* (*p* = 0.002; F_2,15_ = 10.27). In both species, Tukey’s HSD test showed that trees in twelve-species plots did not significantly differ from those in monoculture. There were no significant differences at all among treatments in *Q. ellipsoidalis* (*p* = 0.294; F_2,15_ = 1.330) or *T. americana* (*p* = 0.359; F_2,15_ = 1.099). Considering area-based assimilation, we see similar results, except that there are no significant differences among treatments in *B. papyrifera* (*p* = 0.0892; F_2,15_ = 2.852), only in *A. rubrum*.

#### Instantaneous chlorophyll fluorescence and spectral reflectance

Among all eight species except *B. papyrifera*, dark-acclimated F_v_/F_m_ declined as relative light availability (RLA) increased (Fig. 4). Extracting the species-specific slopes of this relationship, we found that species with high shade tolerance (as quantified by Niinemets & Valladares 2006) had the greatest decline in F_v_/F_m_ as RLA increased (Fig. 5; *R*^*2*^ = 0.601; *p* = 0.025; *t*(5) = −3.167). In most species, non-photochemical quenching (qN) rose with RLA, but in *T. americana* and *A. negundo*, qN was nearly constant across light environments (Fig. 4). Consequently, we found that shade-tolerant species also had smaller rises in qN with RLA (Fig. 5; *R*^*2*^ = 0.619; *p* = 0.022; *t*(5) = −3.276). PRI declined as RLA increased, with statistically indistinguishable slopes among species (Fig. 4).

**Fig. 4:**
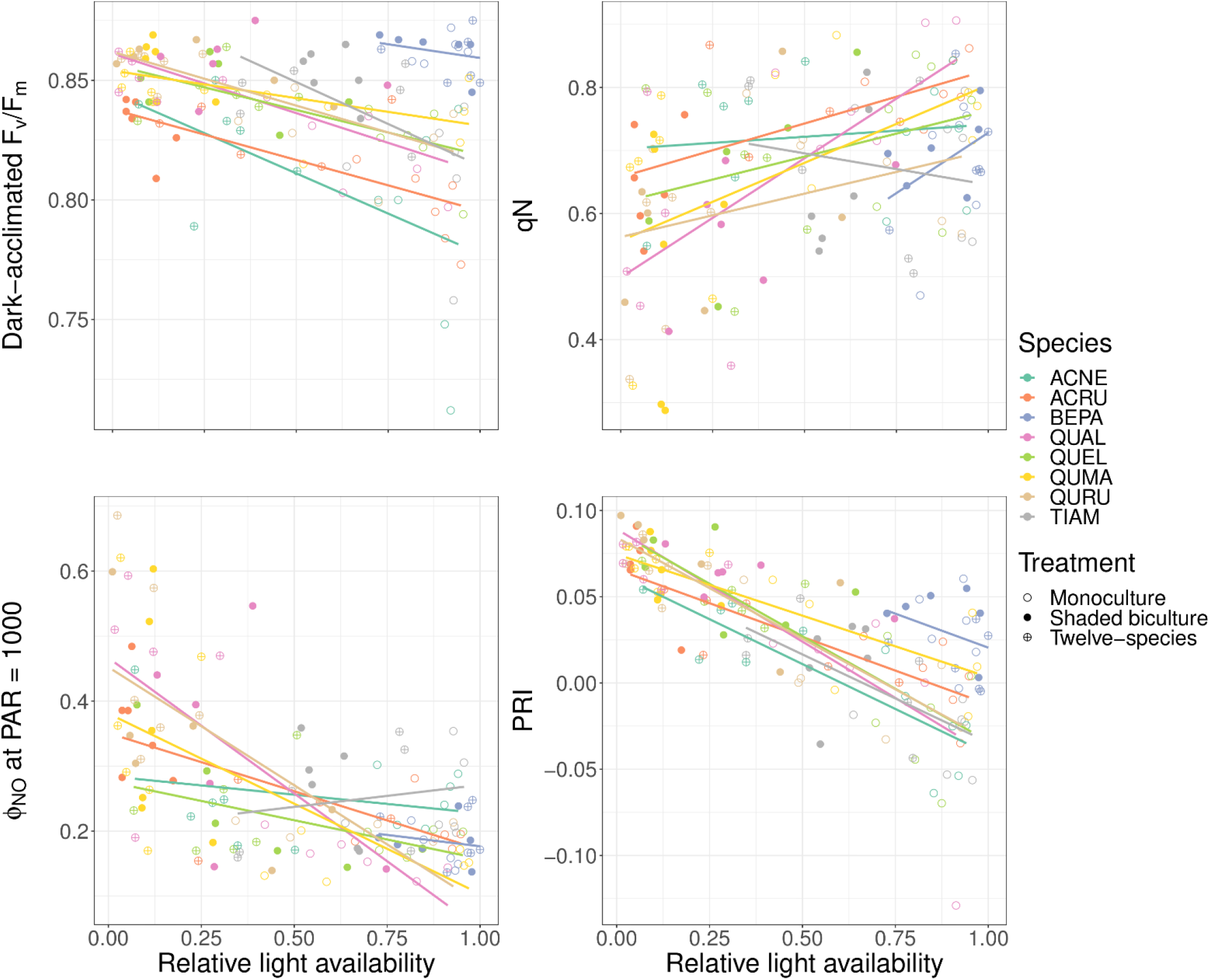
Chlorophyll fluorescence parameters and the photochemical reflectance index (PRI) as a function of relative light availability across species and treatments. Best-fit lines come from species-specific OLS regressions. Dark-acclimated F_v_/F_m_ is the maximal quantum efficiency of PSII photochemistry and declines as a plant becomes more photoinhibited. qN describes a plant’s capacity to use non-photochemical quenching to dissipate energy. ϕ_NO_ represents the proportion of light dissipated in an unregulated way, which may contribute to photodamage. PRI correlates negatively with carotenoid: chlorophyll ratios.

**Fig. 5:**
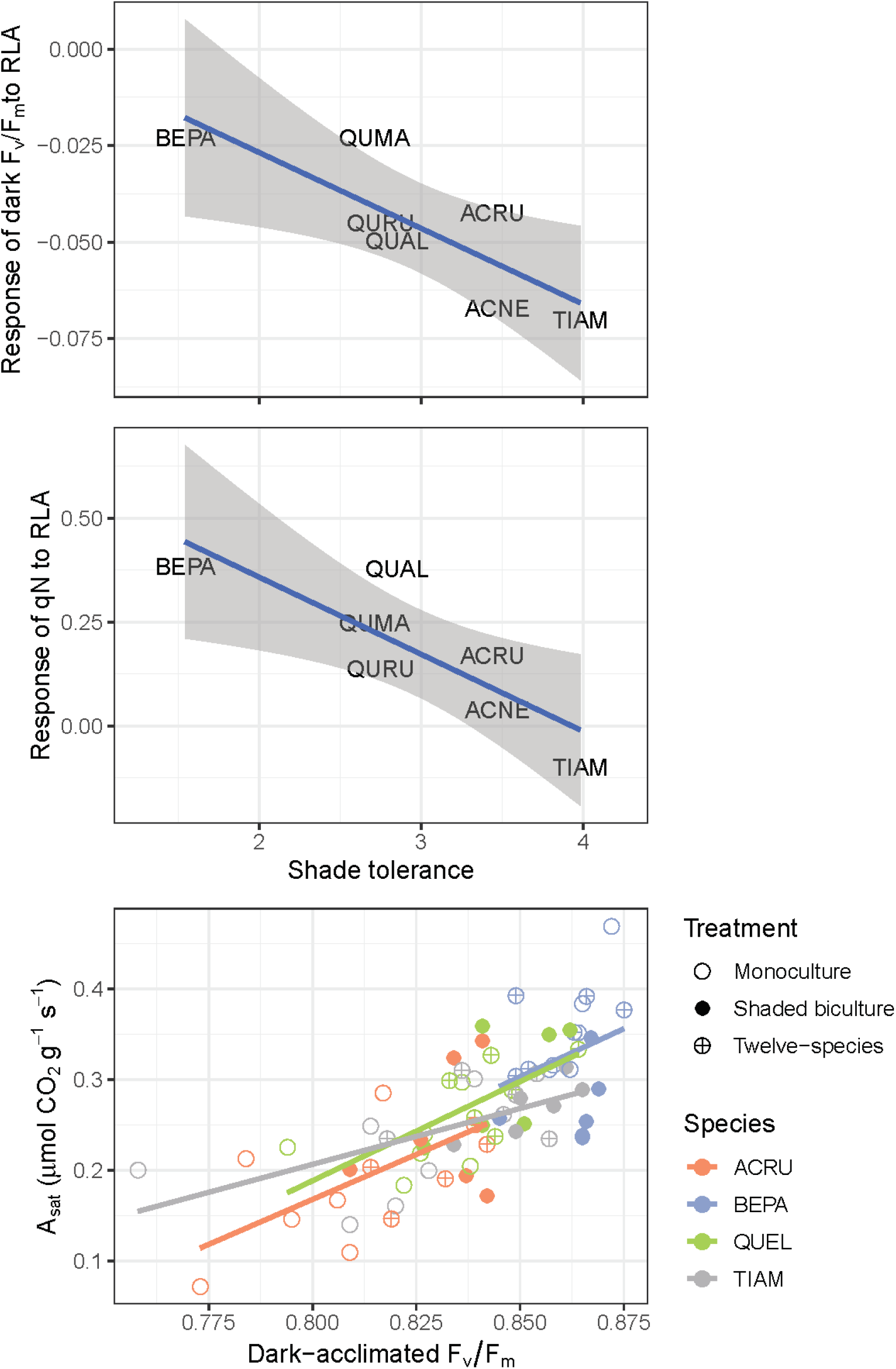
Compared to shade-intolerant species, shade-tolerant species show greater declines in dark-acclimated F_v_/F_m_ (top) and smaller increases in qN (middle) as light availability increases. *A*_*sat*_ derived from light-response curves correlates positively and strongly with dark-acclimated F_v_/F_m_; slopes are statistically indistinguishable among species (bottom). Species codes are found in Table 1.

To see why shade-tolerant species showed such large increases in photoinhibition with light, consider ϕ_PSII_, ϕ_NPQ_, and ϕ_NO_—the quantum yields of PSII photochemistry, non-photochemical quenching, and non-regulated dissipation. In most species, ϕ_NO_ decreased with light in the growth environment (Fig. 4)—for trees growing in the open (RLA ≈ 1), the sum ϕ_PSII_ + ϕ_NPQ_ at 1000 μmol photons m^−2^ s^−1^ was usually above 0.8, keeping ϕ_NO_ close to the oft-cited theoretical minimum of ~0.17 (Tietz et al. 2017; Fig. S7). But in *T. americana* and *A. negundo*, ϕ_NO_ was nearly equal across the gradient of light availability.

To illustrate how photoinhibition affected photosynthetic function, we examined the relationship between dark-acclimated F_v_/F_m_ and *A*_*sat*_. Among the four focal species, dark-acclimated F_v_/F_m_ was positively correlated with mass-based *A*_*sat*_ (OLS regression; *R*^*2*^ = 0.493; *p* < 10^−11^; *t*(69) = 8.304; Fig. 5), and slopes were not significantly different among species (ANCOVA, species × dark-acclimated F_v_/F_m_ interaction; *p* = 0.658; F_3,63_ = 0.539). The same overall relationship holds when *A*_*sat*_ is expressed on an area basis (*R*^*2*^ = 0.447; *p* < 10^−9^; *t*(69) = 7.588), although within species, *A. rubrum* and *B. papyrifera* had no significant relationship.

One way of disentangling the role of photoinhibition from leaf plasticity in influencing photosynthetic rates is to consider the residuals of the species-specific relationships between area-based *A*_*sat*_ and LMA. These residuals quantify how much greater or lesser *A*_*sat*_ is than predicted based on LMA alone for a given species. Dark-acclimated F_v_/F_m_ showed a positive correlation with these residuals in *Q. ellipsoidalis* (*R*^*2*^ = 0.351; *p* = 0.006; *t*(16) = 3.189) and *T. americana* (*R*^*2*^ = 0.368; *p* = 0.004; *t*(16) = 3.304). In these two species, a rise of 0.01 in dark-acclimated F_v_/F_m_ would be predicted to cause a rise of 1.41 and 0.85 μmol m^−2^ s^−1^ in carbon assimilation, respectively. We found no such relationship in *B. papyrifera* or in *A. rubrum*, although in *A. rubrum* alone, dark-acclimated F_v_/F_m_ showed a strong negative correlation with LMA (*R*^*2*^ = 0.460; *p* = 0.002; *t*(15) = −3.826). This multicollinearity makes it hard to estimate the true contribution of either variable to area-based *A*_*sat*_, and leaves open the possibility that a potential positive effect of LMA on *A*_*sat*_ is offset by photoinhibition as light availability increases.

### Phenology

In *T. americana*, leaf abscission in monoculture began in early August, and more than half of all leaves had senesced by September 3 (Fig. 6). In shaded bicultures, more than 90% of leaves remained by September 3, and no survey found greater than 50% senescence until October 1. We assigned the senescence date of each leaf as the date of the first survey by which it had senesced; using this response variable, leaves in monoculture senesced 22 days earlier than those in shaded bicultures (t-test; *p* < 10^−11^; *t*(116) = 7.822). In mid-September, dark-acclimated F_v_/F_m_ averaged 0.745 among remaining leaves in shaded biculture, but only 0.642 in monoculture (t-test; *p* = 0.061; *t*(33) = 1.936).

**Fig. 6:**
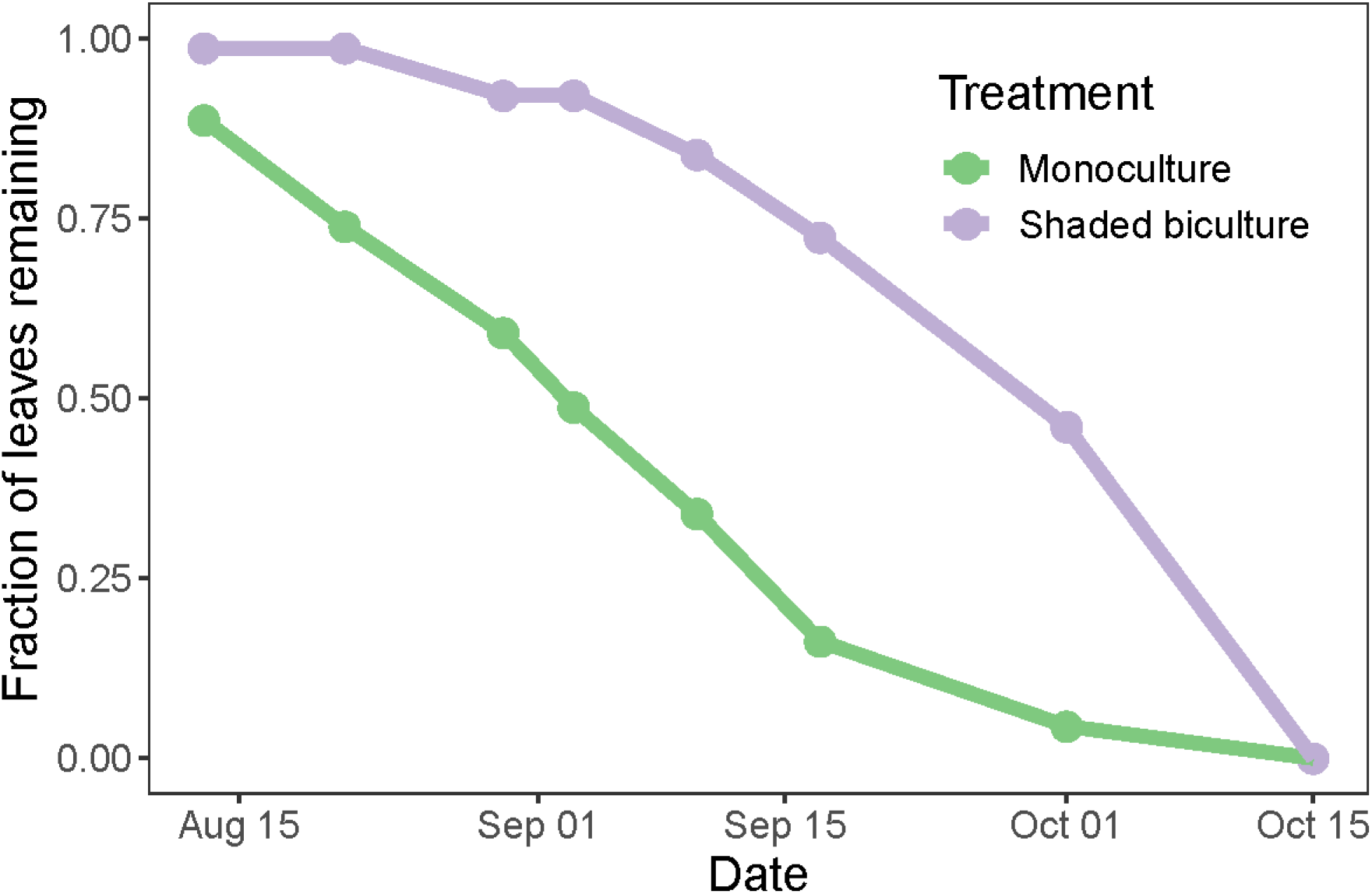
In *Tilia americana*, fall leaf abscission occurs later in monoculture than in shaded biculture.

Our one-time measurement of *A. rubrum* leaves in early October confirmed that this pattern is not limited to *T. americana*. *A. rubrum* trees in shaded biculture plots retained about 56% of their leaves, while those in monoculture retained only 10% (t-test; *p* < 10^−4^; *t*(44) = −4.486).

### Water potential

We removed two data points, one in each treatment, with Ψ_PD_ more negative than −0.8 MPa on the grounds that they may have been subject to measurement error. Across treatments, the mean Ψ_PD_ and Ψ_MD_ were −0.15 and −1.29 MPa. Ψ_PD_ increased with light availability (*R*_*m*_^*2*^ = 0.272; *p* < 10^−5^; *t*(70.0) = −5.152), making it less negative in monocultures than shaded bicultures by an average of 0.07 MPa (Fig. S8). This result holds even including the outliers (*R*_*m*_^*2*^ = 0.085; *p* = 0.014; *t*(54.3) = −2.555). By midday, the pattern reversed; Ψ_MD_ declined with light availability (*R*_*m*_^*2*^ = 0.346; *p* < 10^−4^; *t*(36.6) = 4.450), making it more negative in monocultures than shaded bicultures by an average of 0.33 MPa.

### Leaf angles

Among each of the three species whose leaf angles we measured, leaves were nearer to horizontal in the shaded biculture treatment (Fig. 7). The size of this fixed effect varied among species: 11.0° in *Q. ellipsoidalis* (*p* = 0.005; *t*(34.2) = −3.010), 21.2° in *A. rubrum* (*p* < 10^−9^; *t*(108.4) = −6.807), and 14.3° in *T. americana* (*p* = 0.003; *t*(124.5) = −3.040). Because *T. americana* leaves usually drooped downward in monoculture, leaf angles became less negative in the shade. In *Q. ellipsoidalis*, whose leaves were often inclined upward in monoculture, leaf angles became less positive in the shade. In *A. rubrum*, leaf angles were inclined both up and down in monoculture, but varied less in either direction in the shade.

**Fig. 7:**
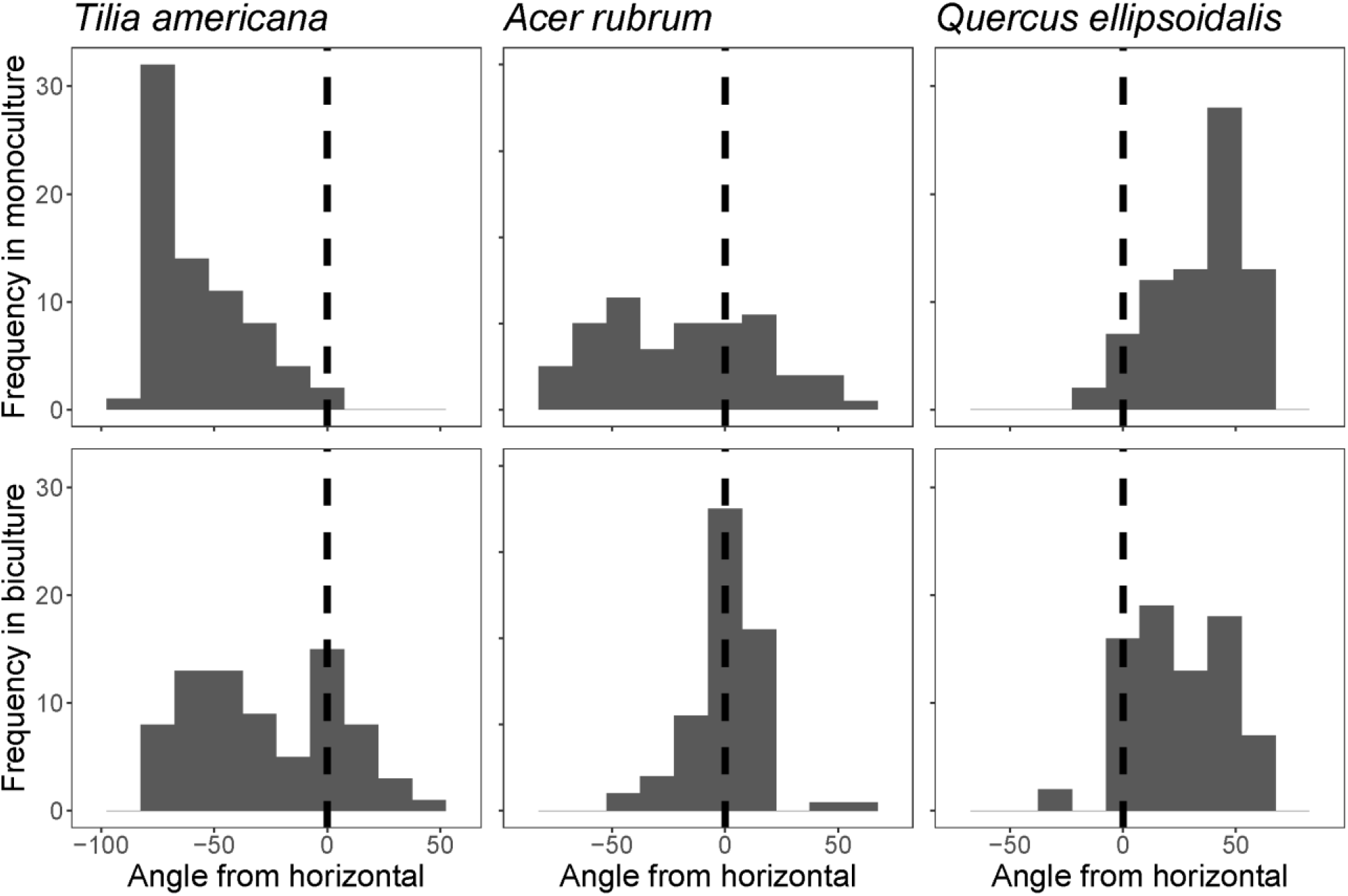
The angles of top leaves of three species are more horizontal in shaded biculture plots than in monoculture.

### Environmental factors

In our survey, soil moisture varied among treatments (ANOVA; *p* < 0.01; F_2,29_ = 6.115); it was lowest (5.7%) in shaded biculture plots and highest (7.2%) in broadleaf monoculture plots. We found that daytime air temperature in the open was higher than in shaded bicultures or twelve-species plots at 62.5% of hourly log-points during the nine-day logging period. The air temperature in monocultures was higher on average by 1.03 °C over this period.

## Discussion

We investigated how species interactions in a tree diversity experiment might emerge from the physiological responses of trees to the light environment created by their neighbors. We found that as neighbor size increased, five species (*A. rubrum*, *Q. alba*, *Q. ellipsoidalis*, *Q. macrocarpa*, and *Q. rubra*) had faster stem growth on average, while the remaining three species (*A. negundo*, *T. americana*, and *B. papyrifera*) had slower stem growth. The five species with declining stem growth responses to neighbor size were shade-intolerant, resistant to photoinhibition in high light, and tended to have lower carbon assimilation in shaded bicultures. Two of the three species with increasing responses were shade-tolerant and susceptible to photoinhibition. The divergent responses of these two groups of species seem driven in part by their tolerance to excess light. The remaining species, *B. papyrifera*, had a positive response to neighbor size despite lower photosynthetic rates. Here, we interpret these patterns further.

### Competition and shade-intolerant species

Individuals of all four *Quercus* species and *A. rubrum* had lower stem growth when surrounded by larger neighbors (Fig. 2). This growth pattern implies that neighborhood interactions were dominated by competition, potentially both above- and belowground. (The fact that soil moisture was lowest in shaded biculture plots suggests an important role for belowground competition.)

The physiological data can help us make sense of these growth patterns. We measured light-response curves of two species in this group: *A. rubrum* and *Q. ellipsoidalis*. In the untransformed chamber light-response curves, both had lower mass-based assimilation rates in monoculture than in the other two treatments (Fig. 3, left). But on an area basis, *A. rubrum* had its lowest assimilation rates in the shaded biculture treatments, and *Q. ellipsoidalis* was similar across all three treatments. The discrepancy between area- and mass-based results is explained by variation in LMA across treatments (Fig. S2).

Multiple structural and physiological factors contribute to explaining variation in photosynthesis across treatments. *Q. ellipsoidalis* and *T. americana* had positive relationships between LMA and area-based *A*_*sat*_, which may imply that much of the variation in LMA is driven by the mass of photosynthetic tissue (Osnas et al. 2018). In contrast, *B. papyrifera* and *A. rubrum* had no correlation between LMA and area-based *A*_*sat*_, which may imply that much of the variation in LMA could be driven by the mass of non-photosynthetic structural tissue. But for *A. rubrum* in particular, there was a strong negative correlation between LMA and dark-acclimated F_v_/F_m_, two variables jointly affected by light availability (Fig. S2; Fig. 4). As a result, a potential positive relationship between LMA and *A*_*sat*_ in *A. rubrum* could have been masked by photoinhibition. We also found evidence that photoinhibition lowers *A*_*sat*_ in *Q. ellipsoidalis* and *T. americana*. Since photoinhibition occurs through damage or downregulation of PSII, it is usually thought to depress photosynthesis by lowering ETR. But both ETR and stomatal conductance mirrored assimilation across treatments in chamber light-response curves (Fig. S5), so both may have some role in explaining why most species had lower assimilation in monoculture at a given chamber light level.

Rescaling the light-response curves allowed us to visualize the costs of shading—the fact that some trees receive less light and face more frequent photosynthetic light limitation than others. Compared to monoculture, *A. rubrum* and *Q. ellipsoidalis* in shaded biculture had lower mass-based carbon assimilation under low light at the top of the canopy, but higher mass-based carbon assimilation under high light (Fig. 3, right). For these two species, shading often appears severe enough to cause light limitation of growth and photosynthesis, especially under otherwise low light.

Given that light availability fluctuates—diurnally, seasonally, and with cloud cover—the treatment that performs best may also vary from moment to moment. We used a time-series of solar radiation to estimate total carbon assimilation throughout July. We found that *A. rubrum* and (only on a mass basis) *B. papyrifera* had lower carbon assimilation in shaded biculture than in the other treatments (Fig. S6). PPFD was under 500 μmol m^−2^ s^−1^ more than 60% of the time during the month; it is under such dim conditions that trees in monoculture have the greatest photosynthetic advantage over those in shaded biculture, no matter what benefits shading may confer at sunnier times (Fig. 3; Fig. S4). Trees in twelve-species plots had similar assimilation rates to those in monoculture in all species. The intermediate light environment in twelve-species plots may have allowed trees to avoid photoinhibition without becoming frequently light-limited (Fig. S2), although we caution that we only measured top leaves, so we cannot directly compare whole-plant carbon gain.

Trees in FAB and elsewhere often vary in size by orders of magnitude, which foregrounds the size-asymmetry of light competition. Research on light competition in biodiversity-ecosystem function research underscores a tension between competitive imbalance caused by size-asymmetry and competitive relaxation caused by light-use complementarity, in which species may partition their exploitation of the light environment (Yachi & Loreau 2007; Sapijanskas et al. 2014; Williams et al. 2017). We leave the potential for light-use complementarity unaddressed, but we show that competitive imbalance can suppress the photosynthesis and growth of species that are poorly adapted to shade. Such imbalances could promote selection effects (*sensu* Loreau & Hector 2001), in which the most productive species become even more dominant in mixtures.

### Facilitation and shade-tolerant species

In *T. americana*, *A. negundo*, and *B. papyrifera*, having larger neighbors increases stem growth (Fig. 2; Fig. S3). The former two are the most shade-tolerant broadleaf species in the experiment, while the last is the most shade-intolerant. Moreover, these three species showed divergent photosynthetic responses to the light environment, which suggests that different mechanisms may explain the positive stem growth response in *B. papyrifera* than in the other two species.

We first discuss the positive trend in *T. americana* and *A. negundo* before returning to *B. papyrifera* in the following section. In these two species, we interpret this trend mainly as a result of facilitation, reinforcing prior results that showed that shade-tolerant species may often have facilitative responses to being shaded (Montgomery et al. 2010). In particular, these two species’ growth trends are best fit by a model where neighbor size is log-transformed (Fig. 2, Appendix 2). Increasing neighbor size thus has positive but diminishing marginal benefits. This outcome may result if neighbor size has a non-linear influence on light availability, or if deep shade has escalating costs that begin to offset its benefits.

Here, the physiological data may help us explain the growth patterns. *T. americana* was among the four focal species, and the only one of the four that had the lowest carbon assimilation rates in the monoculture treatment across most light levels in both chamber and rescaled light-response curves (Fig. 3). The steep decline in dark-acclimated F_v_/F_m_ with increasing light (Fig. 4) suggests that the low *A*_*sat*_ in monoculture results from photoinhibition (Fig. 5). This pattern aligns with *T. americana*’s role as a late-successional dominant species throughout much of its range (Braun 1950). Our finding that carbon gain in *T. americana* can increase under shade is reinforced by Carter and Cavaleri (2018), who found that its assimilation rates may increase down vertical gradients from the upper canopy to the sub-canopy. Nevertheless, we found no significant differences among treatments in estimated assimilation across the whole month of July (Fig. S6), mainly because under low light, monocultures still had higher assimilation rates in the rescaled light-response curves. The amelioration of photoinhibition is very likely to contribute to the positive effect of neighbor size on growth in *T. americana*, but it cannot fully explain it.

We next discuss two mechanisms of facilitation that could also contribute to positive growth responses to neighbor size, but which analyses based on light-response curves do not account for: Delayed senescence and microclimatic effects. Both may contribute to potential positive effects of shading by neighbors in the species that show such responses.

*T. americana* in monoculture dropped its leaves nearly a month earlier into the fall compared to the shaded biculture (Fig. 6). We also found that *A. rubrum* in monoculture had earlier leaf senescence. Given that leaves remained photosynthetically active until shortly before abscission (as suggested by Mattila et al. 2018), shade could allow these trees a longer period of carbon gain. This pattern admits at least two explanations: (1) that plants in the sun are earlier to accumulate phenological cues for senescence related to temperature or photoperiod; or (2) that abiotic stress, especially light and water stress, accelerate senescence (Cavender-Bares et al. 2000; Estiarte & Peñuelas 2015; Brelsford et al. 2019), especially as the temperature declines, rendering leaves more susceptible to photodamage (Juvany et al. 2013; Renner & Zohner 2019). Contrary to the first explanation, air temperatures in full sun were higher than in shaded plots, suggesting that early leaf drop in monoculture was caused by stress. Without measuring the timing of senescence in other species, we do not yet know how general this effect is.

Although we emphasize the role of light stress, other microclimatic factors could contribute to variation in assimilation rates among treatments. For example, an alternate perspective on photoinhibition is that it mainly results not from photodamage but from carbon sink limitation, which leads plants to downregulate light capture and electron transport as a photoprotective response (Adams III et al. 2013). From this perspective, photoinhibition seldom causes carbon limitation or reduces growth rates in nature; rather, slow growth rates cause photoinhibition. A potential cause of sink limitation that could affect monocultures of *T. americana* and *A. negundo* more than shaded bicultures is water limitation, which could physically constrain the expansion of tissues (Tardieu et al. 2014). We found that leaf Ψ_PD_ was slightly more negative in shaded bicultures, while Ψ_MD_ was more negative in monocultures (Fig. S8). (These results are consistent with the idea that soil moisture is higher in low biomass plots, as we found, while daytime VPD is lower in high biomass plots, as Wright et al. (2018) found on hot, dry days at Cedar Creek.) If Ψ_MD_ responds in other tissues as in leaves, highly negative values could arrest growth in monoculture, causing carbohydrates to accumulate in leaves under high light.

But from this perspective, it is hard to explain why this sink limitation would particularly matter in *T. americana* and *B. papyrifera* among the four focal species; for example, *A. rubrum* also has more negative Ψ_MD_ in monoculture (Fig. S8). Moreover, trends in ϕ_NO_ suggest that *T. americana* and *A. negundo* in high light are especially vulnerable to photodamage (Fig. 4). Photoinhibition due to source-sink imbalance is usually accompanied by compensatory upregulation in protective non-photochemical quenching (Adams III et al. 2013), but these two species fail to increase qN under high light (Fig. 4). These findings reinforce the idea that damage, not just downregulation, contributes to the steep declines in dark-acclimated F_v_/F_m_ these species show under increasing light. Sink limitation may contribute to the photoinhibition and the growth patterns we observe in these species, but we believe it is not the main explanation.

The microclimate could also alter photosynthesis in ways our light-response measurements did not capture because we controlled the chamber microclimate. Our water potential measurements show that water stress was not very severe in any treatment, but even modest differences could change the timing of stomatal closure (Brodribb et al. 2003). It also seems plausible that greater light exposure may make leaf temperature higher in monoculture (Schymanski et al. 2013), which could push leaves above thermal optima for photosynthesis. Such microclimatic factors could contribute to the physiological benefits of shading beyond what we could measure in light-response curves.

### Shade avoidance and early-successional species

In the only early-successional species, *B. papyrifera*, the stem growth response is weak and dependent on the model specification (Supplemental Materials, Appendix 2), but it does appear to reflect a longer-term trend that has caused trees with larger neighbors to show faster stem growth in absolute terms. Consider the absolute growth rate from 2017 to 2018: *B. papyrifera* stems grew faster in shaded bicultures (median: 578 cm^3^; 25^th^–75^th^ percentile: 231–1168 cm^3^) than in bicultures with other broadleaf species (median: 496 cm^3^; 25^th^–75^th^ percentile: 87–836 cm^3^). But there is a striking mismatch between the growth patterns and photosynthetic physiology: In untransformed and rescaled light-response curves alike, *B. papyrifera* had lower photosynthetic rates in the shaded biculture compared to the other treatments. Because it tended to overtop even its largest neighbors, the top leaves received nearly the full amount of available light in all treatments (Fig. S2) and showed few signs of photoinhibition (Fig. 4). These results suggest that the increase in stem volume is unrelated to photosynthetic physiology.

One possibility is that rather than (or in addition to) representing an increase in total growth due to facilitation, the positive trend in stem volume in *B. papyrifera* is driven by a competitive shade avoidance response, which could increase allocation to shoot biomass at the expense of root biomass. Such responses tend to be especially strong in shade-intolerant, early-successional species like *B. papyrifera* (Henry & Aarssen 1997; Gilbert et al. 2001). While we could not test directly for a change in the root-to-shoot ratio, we did find evidence that *B. papyrifera*, but not *A. negundo* or *T. americana*, had another symptom of the shade avoidance syndrome: greater allocation to elongation growth near larger neighbors, which caused trees to have smaller diameters than expected based on their height (Henry & Aarssen 1999). This finding may suggest that a broader shade avoidance syndrome may contribute to the positive influence of neighbor size on stem volume in *B. papyrifera*. This finding would be in contrast to another tree diversity experiment, where both root-to-shoot ratios and the allometric relationship between basal area and biomass were nearly unaffected by the local neighborhood (Guillemot et al. 2020). In the absence of direct evidence, changes in root-to-shoot ratios driven by shade avoidance remain a tentative but plausible contributor to positive growth responses in *B. papyrifera*. Such changes in allocation are not mutually exclusive with facilitation, but could obviate the need to invoke facilitation.

### Mechanisms of photoprotection

Plants’ strategies to avoid damage from the stress of excess light can be classified broadly into biochemical and structural strategies. We show that trees in full sun allocate more to biochemical photoprotection, as indicated by lower PRI in all species and higher qN in all species but *T. americana* and *A. negundo* (Fig. 4). Our leaf angle survey is also consistent with the idea that plants may steeply incline their leaves to avoid intercepting excess light—a form of structural photoprotection (Fig. 6). Leaves that are steeply inclined intercept less light per unit area, particularly during midday, when solar radiation is otherwise most intense. The dramatic trend in leaf angles suggests that gross structural characteristics allow these species use to regulate their light absorption.

Some plants may lack perfect photoprotective mechanisms in part because they are adapted to environments where the costs of high photoprotective investment outweigh the benefits. Delayed relaxation of non-photochemical quenching may hinder photosynthesis even when light declines enough to relieve the imminent threat of damage (Zhu et al. 2004; Murchie & Niyogi 2011; Kromdijk et al. 2016). Building pigments and proteins for photoprotection uses resources that plants could otherwise allocate to chlorophyll, RuBisCO, or structural tissue components, although the concentrations of such photoprotective compounds are often low. Steep leaf angles may also decrease photosynthesis by reducing light interception even when light is limiting—and all plants experience light-limiting conditions at least sometimes, as during cloudy days or nighttime. All the same, plants in more open environments may need photoprotective mechanisms to avoid even greater costs due to photodamage under brighter conditions. That most species upregulate photoprotection in high light, despite the costs, implies that these species may otherwise incur a risk of damage from excess light. These findings reinforce that light stress and its alleviation through shading can contribute to net species interactions.

### Competition and facilitation in biodiversity-ecosystem function relationships

The finding that shade can increase growth and photosynthesis has precedents. Ball et al. (1991) and Egerton et al. (2000) showed that shade enhances assimilation and growth of evergreen *Eucalyptus* seedlings during the winter, when cold may cause photoinhibition. Howell et al. (2002) also found that among evergreen divaricating shrubs of New Zealand, leafless outer branches increase photosynthesis during the winter by reducing photoinhibition. Species can also facilitate each other from light stress in communities of phytoplankton (Gerla et al. 2011), which nearly all face some risk of photoinhibition (Edwards et al. 2015; Edwards et al. 2016) These studies show that facilitation through shading can occur in many environments, and perhaps more so in otherwise stressful ones.

Much of the work on facilitation in plant communities is guided by the stress-gradient hypothesis (SGH), which proposes that facilitative interactions are more common in stressful environments (Bertness & Callaway 1994; Maestre et al. 2009). We can place photoinhibition in light of the SGH as follows: Stressful conditions limit the use of light for photochemistry, turning high light exposure into an additional stressor. Under such stressful conditions, shade from neighbors can ameliorate stress, resulting in facilitation. Under benign conditions, where plants can use more of the available light for photochemistry, high light exposure may not cause damage. Here, shading by neighbors is more likely to cause light limitation, resulting in an adverse, competitive effect on growth. As we show here, whether a species is stressed by its environment also depends on its own physiological tolerances.

The SGH brought attention to facilitation in plant ecology, including in biodiversity-ecosystem function research. For example, Mulder et al. (2001) showed that in experimental moss communities, biodiversity increased productivity only under drought, as species interactions became more positive. But the mechanisms and consequences of facilitation are still overlooked in biodiversity-ecosystem function research (Wright et al. 2017). A small but growing literature based in biodiversity experiments has investigated various mechanisms through which plant species can facilitate each other, including amelioration of water stress (Wright et al. 2015), dilution of natural enemies (Grossman et al. 2019), nitrogen fixation by legume-rhizobia mutualisms (Temperton et al. 2007), and other forms of nutrient-related belowground facilitation (Li et al. 2014). We show that shading can reduce stress induced by the light environment—a specific form of abiotic stress amelioration, in which certain species alter the local environment such that it is less stressful to other species (Wright et al. 2017). But shading can also cause light limitation, resulting in a net competitive effect.

During the first three years of the FAB experiment, Grossman et al. (2017) found that diverse plots overyield—they are more productive than a weighted average of the constituent species in monoculture. Using the partition of Loreau & Hector (2001), most overyielding in FAB was a result of complementarity effects, which are often interpreted as arising from synergistic interactions among species, including facilitation. Overyielding is a hallmark of positive biodiversity-ecosystem function relationships, and a whole community can only overyield if at least one of its constituent species overyields. Even though we test how growth changes with neighbor size, not neighbor diversity, we show that shading allows some species to grow faster in shaded bicultures or twelve-species plots. In the same experiment, Grossman et al. (2017) found that *T. americana* had the greatest overyielding in multi-species plots of any broadleaf species, especially when growing with conifers or *B. papyrifera*. Grossman et al. (2017) further showed that conifer partners in these species combinations also grew faster than they did in monoculture, resulting in community overyielding.

The two broadleaf species that show the strongest evidence for facilitation by shade had low tolerance to excess light, and likely benefited from the milder microclimate their larger neighbors created. The conifer species, which are fast-growing and less shade-tolerant (except for *P. strobus*), may also have grown faster in bicultures with broadleaf species because they faced less competition for light than in monoculture. Other researchers have found that heterogeneity in shade tolerance explains the effect of tree diversity on productivity, but they attributed this result to higher stand-level light capture enabled by crown complementarity (Zhang et al. 2012; Toïgo et al. 2017; Searle and Chen 2019). We show that facilitation of shade-tolerant species by shade-intolerant species is an equally compelling (and perhaps complementary) explanation.

In the literature on biodiversity and ecosystem function, the attempt to parse out mechanisms— such as niche partitioning and facilitation—often depends on statistical partitions of biomass or productivity data (Loreau & Hector 2001; Fox & Kerr 2012) whose interpretation is often debated (Carroll et al. 2011; Pillai & Gouhier 2019). But because productivity has physiological determinants, we can explain its patterns using physiological measurements. In this study, survey-based tree growth data and physiological measurements both show that certain species respond positively and others respond negatively to the size of their neighbors. But only the physiological measurements allow us to explain this divergence—they show that these interactions emerge in part from the photosynthetic responses of trees to the light environment created by their neighbors. The insights gained from such physiological techniques may benefit plant ecology more widely as we seek to explain community patterns in terms of basic aspects of plant function.

## Acknowledgements

University of Minnesota, including Cedar Creek ESR, lies on the ancestral, traditional, and contemporary Land of the Dakota. Conversations with Jake Grossman and Laura Williams inspired much of this research. Beth Fallon, German Vargas G., Artur Stefanski, Daniel Stanton, and Danielle Way all gave valuable advice about measuring photosynthesis. We are indebted to Cathleen Nguyen, Chris Buyarski, Troy Mielke, Kally Worm, Jim Krueger, Pam Barnes, Mark Saxhaug, Susan Barrott, and Dan Bahauddin for making research in FAB possible. Sarah Hobbie and Peter Reich helped design the FAB experiment, and countless Cedar Creek interns have taken part in the stem growth survey. The Cavender-Bares lab (especially Jake Grossman and Gerard Sapès), Daniel Stanton, German Vargas G., Artur Stefanski, David Kramer, and the UMN Physiological Ecology Group all provided feedback on the results or manuscript. FAB is maintained with support from National Science Foundation under DEB #1234162 to Cedar Creek LTER. Spectral measurements were conducted as part of NSF/NASA DEB #1342778 to J.C.B. and R.M. S.K. was supported by an NSF Graduate Research Fellowship (Grant No. 00039202) and a UMN Doctoral Dissertation Fellowship. R.M. is also supported by Minnesota Agricultural Experiment Station project MIN-42-060.

## Supplement

## Appendix 1

## Photosynthetic light-response curve methods

We measured photosynthetic light-response curves from the four focal species (*Acer negundo*, *Betula papyrifera*, *Quercus ellipsoidalis*, and *Tilia americana*) during July 2018 using an LI-6400 gas exchange system (LI-COR BioSciences, Lincoln, NE, USA) with a 6400-40 Leaf Chamber Fluorometer head. From each tree (*n* = 72 total), we selected a fully expanded, sun-exposed leaf with no visible disease or herbivory. We noted the angle of each leaf relative to horizontal to the nearest 15° before beginning measurements. Each curve had nine steps in descending order of brightness: 2000, 1500, 1000, 500, 200, 100, 50, 20, and 0 μmol m^−2^ s ^−1^.

During each curve, we controlled environmental conditions to reduce the immediate effects of environmental conditions other than light on stomatal conductance. We kept relative humidity at a favorable level, typically between 65 and 75%, and set the block temperature as close as possible to 25 °C. The reference CO_2_ concentration remained at 400±10 ppm. We kept each leaf in the chamber for at least five minutes before beginning each curve. We checked that outputs were stable, and matched IRGAs before each measurement.

Since it is hard to control chamber conditions perfectly, any differences that remain among treatments could contribute to the variation we observe in photosynthetic function. During light-response curves, we found no difference among treatments in leaf-to-air VPD within the chamber. There were significant but slight differences in leaf temperature inside the chamber; leaves in monoculture, shaded biculture, and twelve-species plots had average temperatures of 27.7 °C, 26.5 °C, and 27.0 °C, respectively. Previous studies on the temperature response of some of the same species in Minnesota have typically found thermal optima below 26 °C (Sendall et al. 2015). But while cooler leaves may be closer to thermal optima, the small differences we find in leaf temperature cannot explain much of the variation in assimilation rates.

In all but five light-response curves, we also used a saturating pulse to measure light-acclimated chlorophyll fluorescence at each light level. These data allowed us to calculate ϕ_PSII_, a measure of the efficiency of Photosystem II (PSII) at each actinic (or photosynthetic) light level, as:

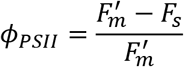

where *F*_*s*_ is the steady-state fluorescence yield under actinic light, and *F*_*m*_’ is the maximum fluorescence yield of the actinic light-acclimated sample following a saturating light pulse, which temporarily closes all PSII reaction centers. From this parameter, we calculated electron transport rate as:

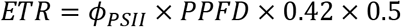

where PPFD is photosynthetic photon flux density (the light level inside the chamber, in μmol m^−2^ s ^−1^), 0.42 is an estimate of leaf absorptance, and 0.5 is an estimate of the fraction of photons that are captured by PSII rather than PSI. Electron transport rate is a key parameter because electron transport is one of the main steps that is capable of limiting photosynthesis (Farquhar et al. 1980), especially when light availability or PSII efficiency are low.

## Chlorophyll fluorescence parameters

As described in the main text, we used dark- and light-acclimated measurements of chlorophyll fluorescence to calculate several derived parameters, from which we make inferences about photosynthetic and photoprotective function. Most importantly, we wanted to assess: (1) capacity for non-photochemical quenching, as measured through the parameters qN or NPQ; and (2) the partitioning of light energy dissipation among pathways, as assessed using the quantum yields ϕ_PSII_, ϕ_NPQ_, and ϕ_NO_.

These derived chlorophyll fluorescence parameters are calculated from a few basic measurements taken during the dark- and light-acclimated measurements, including: *F*_*o*_, the minimum fluorescence yield of a dark-acclimated sample with all PSII reaction centers open; *Fo’*, the minimum fluorescence yield of a light-acclimated sample with all PSII reaction centers open (following far-red light exposure); and *F*_*m*_, the maximum fluorescence yield of a dark-acclimated sample following a saturating pulse. *F*_*v*_, the variable fluorescence yield of a dark-acclimated sample, is calculated as *F*_*m*_ – *F*_*o*_.

The parameters qN and NPQ are meant to indicate the rate constant of the thermal dissipation of energy from PSII (Bilger & Schreiber 1986; Bilger & Björkman 1990). These parameters increase with the use of NPQ to dissipate light—in this case, under exposure to 1000 μmol m^−2^ s ^−1^. They are calculated, respectively, as:

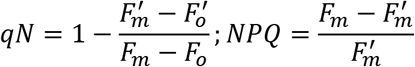

In our dataset, qN is highly correlated (*R*^*2*^ = 0.96) with the natural logarithm of NPQ; we chose to report qN to easily visualize variation among leaves with low NPQ and to help satisfy the error term normality assumption of OLS regression.

The quantum yields ϕ_PSII_, ϕ_NPQ_, and ϕ_NO_ represent the proportions of absorbed light energy that are dissipated through photochemistry, non-photochemical quenching, and non-regulated dissipation. Since these pathways exhaust the options, their quantum yields sum to 1. We calculated them following the ‘lake model’ derivation of Kramer et al. (2004), which treats photosynthetic reaction centers as connected by shared antennae. We achieve qualitatively similar results using the simpler analogues found in Hendricksen et al. (2005).

The parameter ϕ_PSII_ is calculated as shown above (Supplemental Methods, *Photosynthetic light-response curve procedure*). In the derivation from Kramer et al. (2004), ϕ_NPQ_ and ϕ_NO_ are calculated using the parameter qL, an estimate of the fraction of open PSII reaction centers. qL is calculated as:

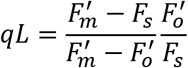

Given this parameter, we can calculate ϕ_NO_ as:

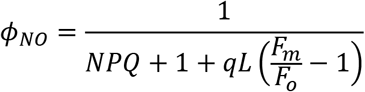

and given the other two quantum yields, we can calculate ϕ_NPQ_ as:

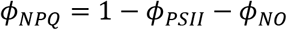

Many researchers, including Murchie and Lawson (2013) and Tietz et al. (2017) treat ~0.83 as a theoretical maximum of ϕ_PSII_ (or dark-acclimated F_v_/F_m_) after a period of dark-acclimation long enough to reduce non-photochemical quenching to zero. Similarly, one may treat ~0.83 as a theoretical maximum of ϕ_PSII_ + ϕ_NPQ_, deviations from which indicate high ϕ_NO_, and thus, a declining ability to regulate energy dissipation. We show this putative maximum in our ternary plot of the quantum yields (Fig. S7).

## Appendix 2

Here, we compare statistical models testing the influence of average neighbor volume on relative growth rate (RGR) of focal individuals (or whole plots). The *t*-statistics, degrees of freedom (df), and *p*-values provided are only for the parameter of interest (*β*_*1*_, the slope term associated with average neighbor volume), even for multivariate models. For the mixed-effect models with plot incorporated as a random intercept, we calculated marginal and conditional *R*^*2*^ using the method of Nakagawa et al. (2017) as implemented in the *R* package *MuMIn v. 1.43.17* (Bartoń 2020), and estimated *p*-values using Satterthwaite’s degrees of freedom method, as implemented in the *R* package *lmerTest v.3.1.2* (Kuznetsova et al. 2017). For the plot-aggregated fixed-effect model, we report a standard adjusted *R*^*2*^. As usual, the *R*^*2*^ values are calculated for the full model, including when models have multiple fixed effects.

In the main text, we present summary statistics from models including only neighbor size as a fixed effect, either untransformed or log-transformed (A or B), based on which fits better for a given species. The results, including slope estimates, are mostly robust to the model specification (C-E). We highlight in blue the results presented in the main text, and in yellow, cases where our inferences about how species respond to neighbor volume would be different under an alternate model specification (based on comparing *p*-values to an α = 0.05 threshold). For models that include focal individual size in 2016, we note that these values are not usually correlated with RGR between 2016 and 2018 (*R*^*2*^ < 0.01 except in *Acer negundo*), so there is no problem of multicollinearity.

The final three models have independent variables other than focal individual RGR. In F, we predict plot-level RGR from the plot-scale average of each individual’s average neighbor biovolume. In G and H, we predict the absolute growth rate of focal individuals, which allow us to compare model parameters including (G) and excluding (H) trees that died between 2016 and 2018. Because of extreme heteroskedasticity and non-normality of errors, we implemented these models using robust regression with a smoothed Huber loss function in *R* package *robustlmm 2.3* (Koller et al. 2016).

| Species | $\beta_I \times 10^4$ (SE $\times 10^4$) | $R_m^2$ | $R_c^2$ | $t$ | df | $p$ |
| --- | --- | --- | --- | --- | --- | --- |
| *Acer negundo* | 0.394 (0.324) | 0.003 | 0.374 | 1.217 | 380 | 0.224 |
| *A. rubrum* | -0.775 (0.238) | 0.027 | 0.191 | -3.257 | 366 | 0.00123 |
| *Betula papyrifera* | 0.279 (0.140) | 0.010 | 0.056 | 1.993 | 124 | 0.049 |
| *Quercus alba* | -1.062 (0.270) | 0.030 | 0.164 | -3.932 | 486 | $9.65 \times 10^{-5}$ |
| *Q. ellipsoidalis* | -1.270 (0.232) | 0.051 | 0.108 | -5.485 | 341 | $8.05 \times 10^{-8}$ |
| *Q. macrocarpa* | -0.423 (0.203) | 0.006 | 0.218 | -2.079 | 697 | 0.0380 |
| *Q. rubra* | -1.405 (0.243) | 0.053 | 0.087 | -5.785 | 444 | $1.37 \times 10^{-8}$ |
| *Tilia americana* | 0.348 (0.191) | 0.006 | 0.138 | 1.818 | 481 | 0.0697 |

B. $\text{FocalRGR}_{2016-2018} = \alpha + \beta_I \times \log(\text{NeighborVolume}) + (1|\text{Plot})$
| Species | $\beta_I \times 10^2$ (SE $\times 10^2$) | $R_m^2$ | $R_c^2$ | $t$ | df | $p$ |
| --- | --- | --- | --- | --- | --- | --- |
| *Acer negundo* | 4.005 (1.697) | 0.014 | 0.364 | 2.361 | 380 | 0.0187 |
| *A. rubrum* | -0.963 (2.293) | 0.001 | 0.182 | -0.420 | 337 | 0.675 |
| *Betula papyrifera* | 2.498 (1.958) | 0.004 | 0.046 | 1.276 | 167 | 0.204 |
| *Quercus alba* | -6.403 (2.372) | 0.018 | 0.164 | -2.700 | 367 | 0.00725 |
| *Q. ellipsoidalis* | -6.292 (1.928) | 0.021 | 0.093 | -3.264 | 282 | 0.00124 |
| *Q. macrocarpa* | -1.742 (1.768) | 0.002 | 0.227 | -0.985 | 661 | 0.325 |
| *Q. rubra* | -11.119 (2.083) | 0.049 | 0.076 | -5.339 | 224 | $2.29 \times 10^{-7}$ |
| *Tilia americana* | 6.680 (2.174) | 0.018 | 0.143 | 3.073 | 405 | 0.00226 |

| Species | $\beta_I \times 10^4$ (SE $\times 10^4$) | $R_m^2$ | $R_c^2$ | $t$ | df | $p$ |
| --- | --- | --- | --- | --- | --- | --- |
| *Acer negundo* | 0.481 (0.332) | 0.005 | 0.356 | 1.447 | 344 | 0.149 |
| *A. rubrum* | -0.830 (0.249) | 0.030 | 0.204 | -3.338 | 359 | $9.32 \times 10^{-4}$ |
| *Betula papyrifera* | 0.264 (0.145) | 0.009 | 0.069 | 1.823 | 136 | 0.0705 |
| *Quercus alba* | -0.985 (0.281) | 0.026 | 0.180 | -3.502 | 448 | $5.09 \times 10^{-4}$ |
| *Q. ellipsoidalis* | -1.217 (0.237) | 0.048 | 0.126 | -5.135 | 372 | $4.56 \times 10^{-7}$ |
| *Q. macrocarpa* | -0.305 (0.206) | 0.003 | 0.241 | -1.480 | 653 | 0.139 |
| *Q. rubra* | -1.381 (0.248) | 0.052 | 0.081 | -5.580 | 407 | $4.40 \times 10^{-8}$ |
| *Tilia americana* | 0.321 (0.195) | 0.005 | 0.132 | 1.651 | 430 | 0.0996 |

| Species | $\beta_I \times 10^4$ (SE $\times 10^4$) | $R_m^2$ | $R_c^2$ | $t$ | df | $p$ |
| --- | --- | --- | --- | --- | --- | --- |
| *Acer negundo* | 0.393 (0.322) | 0.013 | 0.344 | 1.219 | 376 | 0.223 |
| *A. rubrum* | -0.803 (0.242) | 0.027 | 0.187 | -3.322 | 389 | $9.79 \times 10^{-4}$ |
| *Betula papyrifera* | 0.268 (0.140) | 0.011 | 0.056 | 1.912 | 130 | 0.058 |
| *Quercus alba* | -1.045 (0.272) | 0.031 | 0.165 | -3.845 | 484 | $1.37 \times 10^{-4}$ |
| *Q. ellipsoidalis* | -1.270 (0.232) | 0.051 | 0.108 | -5.480 | 344 | $8.26 \times 10^{-8}$ |
| *Q. macrocarpa* | -0.423 (0.203) | 0.008 | 0.225 | -2.080 | 699 | 0.038 |
| *Q. rubra* | -1.398 (0.242) | 0.057 | 0.089 | -5.769 | 435 | $1.52 \times 10^{-8}$ |
| *Tilia americana* | 0.352 (0.192) | 0.007 | 0.143 | 1.839 | 492 | 0.0666 |

| Species | $\beta_1 \times 10^2$ (SE $\times 10^2$) | $R_m^2$ | $R_c^2$ | $t$ | df | $p$ |
| --- | --- | --- | --- | --- | --- | --- |
| *Acer negundo* | 3.851 (1.692) | 0.025 | 0.339 | 2.277 | 377 | 0.0234 |
| *A. rubrum* | -0.983 (2.315) | 0.001 | 0.181 | -0.424 | 358 | 0.672 |
| *Betula papyrifera* | 2.324 (1.970) | 0.005 | 0.046 | 1.179 | 181 | 0.240 |
| *Quercus alba* | -6.246 (2.382) | 0.020 | 0.167 | -2.622 | 372 | 0.00910 |
| *Q. ellipsoidalis* | -6.312 (1.934) | 0.021 | 0.094 | -3.264 | 290 | 0.00123 |
| *Q. macrocarpa* | -1.702 (1.769) | 0.003 | 0.233 | -0.962 | 665 | 0.337 |
| *Q. rubra* | -10.930 (2.081) | 0.052 | 0.078 | -5.250 | 220 | $3.59 \times 10^{-7}$ |
| *Tilia americana* | 6.752 (2.178) | 0.018 | 0.148 | 3.100 | 416 | 0.00206 |

F. $\text{PlotRGR}_{2016-2018} = \alpha + \beta_1 \times \text{MeanPlotNeighborVolume}$
| Species | $\beta_1 \times 10^4$ (SE $\times 10^4$) | $R_{adj}^2$ | $t$ | df | $p$ |
| --- | --- | --- | --- | --- | --- |
| *Acer negundo* | -0.615 (1.339) | -0.037 | -0.459 | 21 | 0.651 |
| *A. rubrum* | -0.589 (0.455) | 0.022 | -1.296 | 29 | 0.205 |
| *Betula papyrifera* | 0.126 (0.302) | -0.030 | 0.417 | 27 | 0.680 |
| *Quercus alba* | -2.258 (0.620) | 0.297 | -3.642 | 28 | 0.00109 |
| *Q. ellipsoidalis* | -0.870 (0.722) | 0.014 | -1.205 | 31 | 0.237 |
| *Q. macrocarpa* | -2.225 (0.629) | 0.265 | -3.540 | 31 | 0.00129 |
| *Q. rubra* | -1.237 (0.695) | 0.070 | -1.781 | 28 | 0.085 |
| *Tilia americana* | 1.026 (0.257) | 0.312 | 3.995 | 32 | $3.55 \times 10^{-4}$ |

G. $\text{FocalAbsoluteGrowth}_{2017-2018} = \alpha + \beta_1 \times \text{NeighborVolume} + (1|\text{Plot})$
| Species | $\beta_1 \times 10^2$ (SE $\times 10^2$) | $R_m^2$ | $R_c^2$ | $t$ |
| --- | --- | --- | --- | --- |
| *Acer negundo* | 0.301 (0.141) | 0.009 | 0.395 | 2.133 |
| *A. rubrum* | -0.482 (0.159) | 0.024 | 0.200 | -3.040 |
| *Betula papyrifera* | 8.023 (2.023) | 0.044 | 0.215 | 3.966 |
| *Quercus alba* | -0.652 (0.240) | 0.014 | 0.057 | -2.715 |
| *Q. ellipsoidalis* | -0.674 (0.201) | 0.018 | 0.041 | -3.359 |
| *Q. macrocarpa* | -0.582 (0.208) | 0.012 | 0.079 | -2.802 |
| *Q. rubra* | -1.807 (0.345) | 0.039 | 0.039 | -5.236 |
| *Tilia americana* | 5.357 (0.918) | 0.057 | 0.130 | 5.838 |

| Species | $\beta_1 \times 10^2$ (SE $\times 10^2$) | $R_m^2$ | $R_c^2$ | $t$ |
| --- | --- | --- | --- | --- |
| *Acer negundo* | 0.315 (0.176) | 0.007 | 0.449 | 1.794 |
| *A. rubrum* | -0.237 (0.180) | 0.005 | 0.212 | -1.319 |
| *Betula papyrifera* | 9.208 (2.027) | 0.059 | 0.254 | 4.543 |
| *Quercus alba* | -0.618 (0.280) | 0.010 | 0.054 | -2.207 |
| *Q. ellipsoidalis* | -0.674 (0.221) | 0.016 | 0.045 | -3.051 |
| *Q. macrocarpa* | -0.610 (0.213) | 0.013 | 0.073 | -2.866 |
| *Q. rubra* | -1.804 (0.384) | 0.033 | 0.033 | -4.699 |
| *Tilia americana* | 6.212 (0.954) | 0.072 | 0.175 | 6.511 |

---

**Fig. S1:**
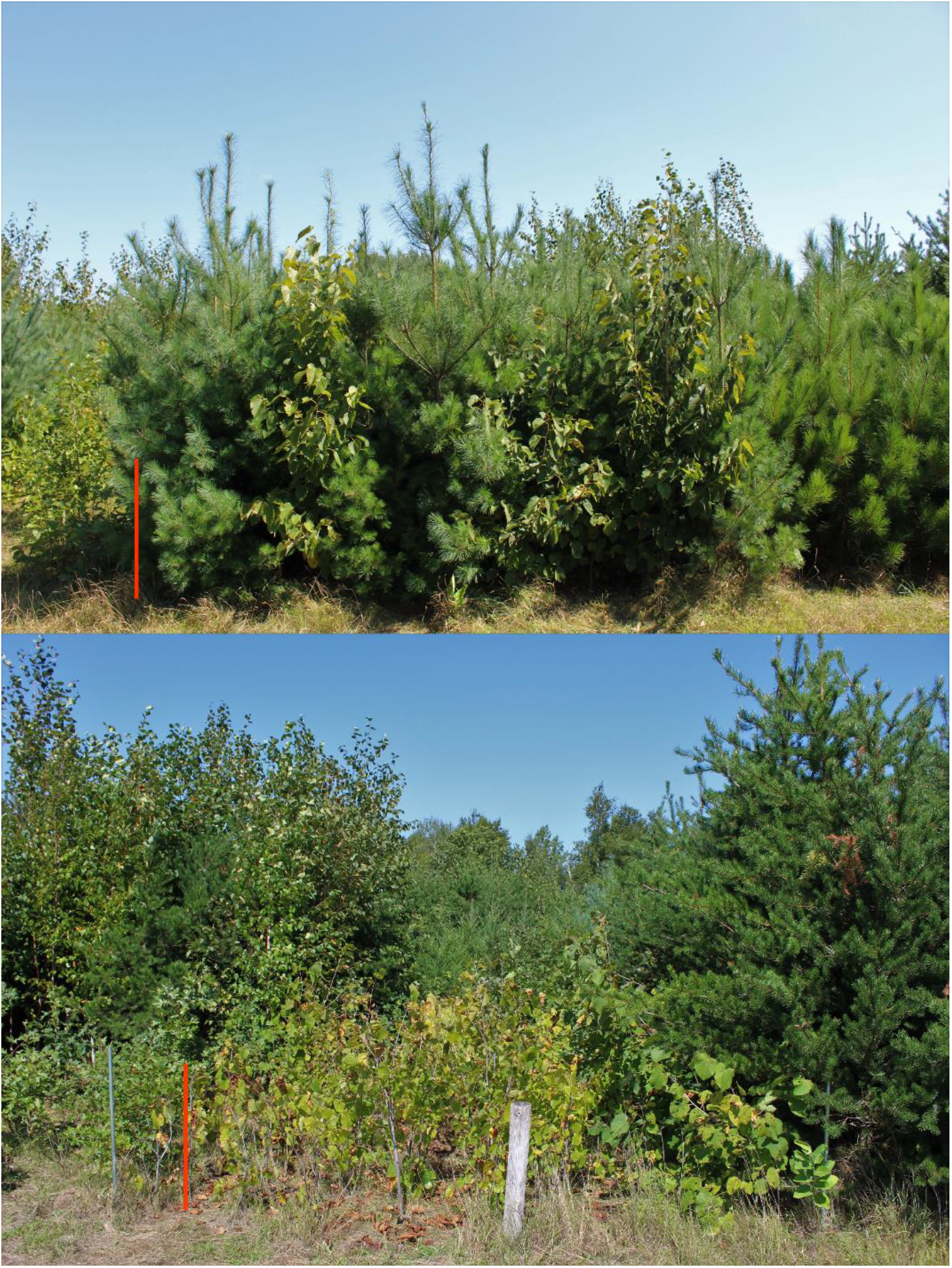
Example images of *Tilia americana* in shaded biculture (top) and monoculture (bottom). Red scale bars represent about 1 m in the foreground.

**Fig. S2:**
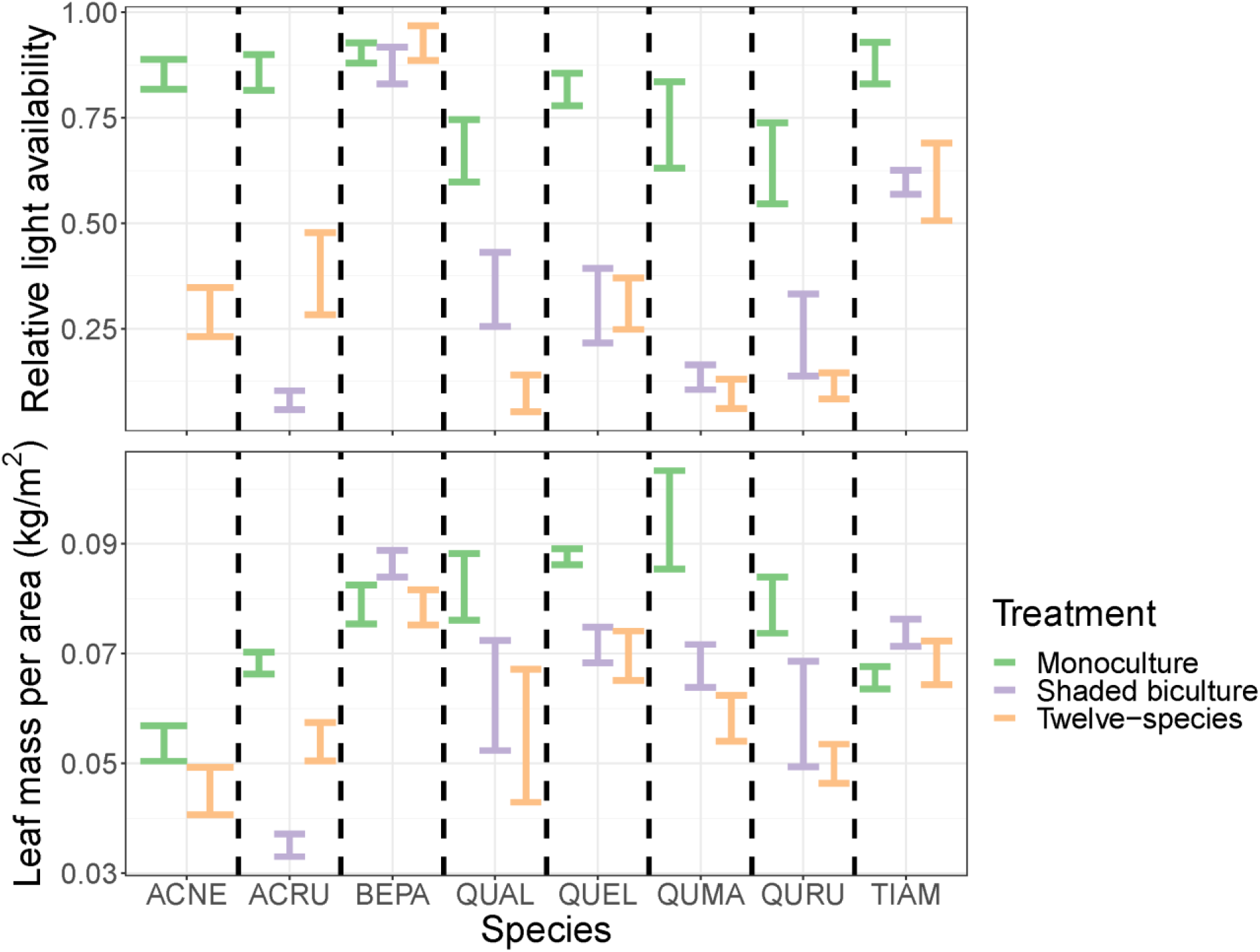
Relative light availability (top) and leaf mass per area (bottom) of each species across treatments. We show LMA values from all leaves used in carbon assimilation and chlorophyll fluorescence measurements. Error bars in left panels are ± 1 SE.

**Fig. S3:**
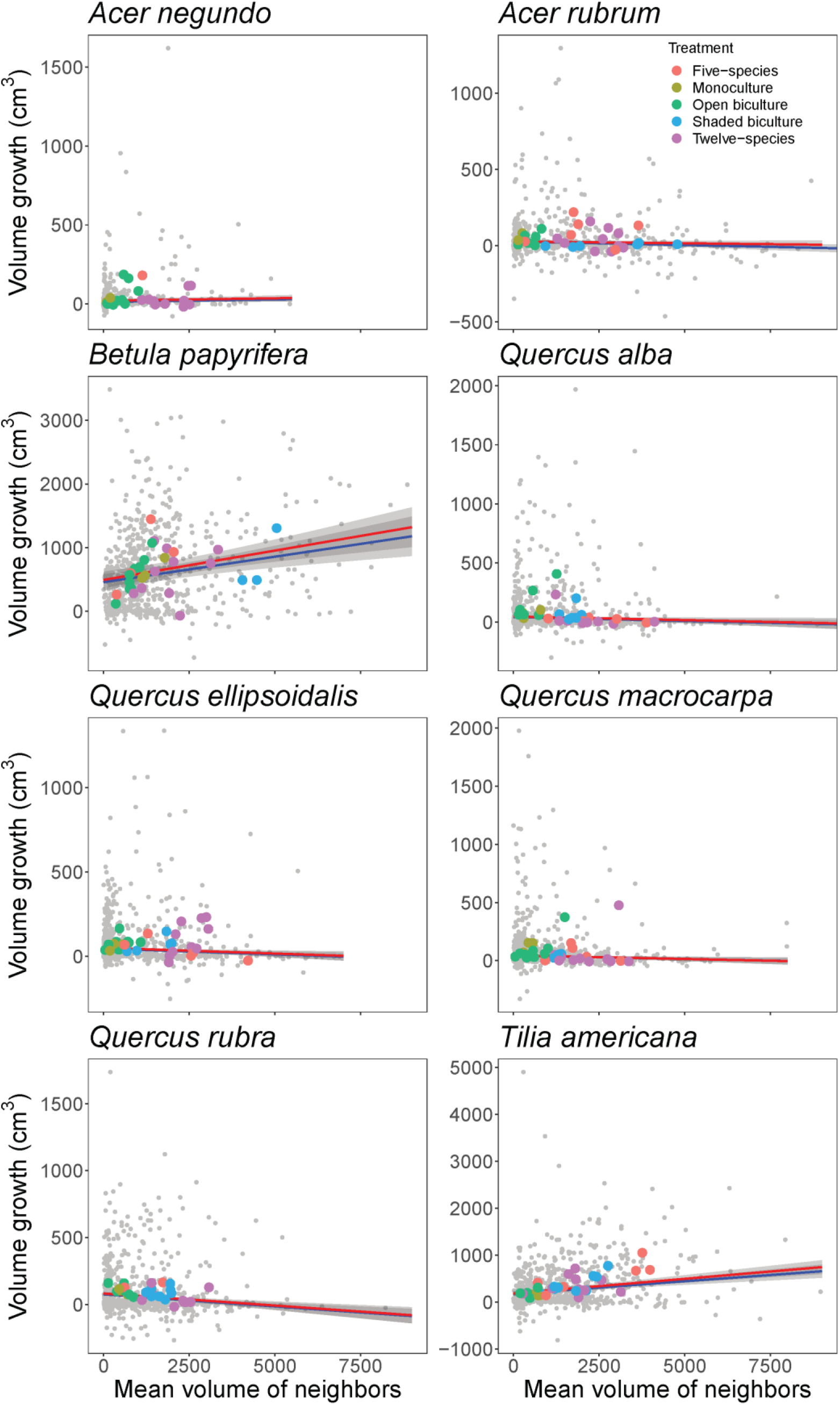
Growth of woody stem volume in individuals of each species between fall 2017 and 2018 as a function of the average stem volume of all neighbors. Gray dots represent individuals, while colored dots are aggregated to the plot scale and color-coded by treatment. Based on mixed-effects robust regression models with plot as a random intercept, two lines are fit for each species: One (blue) to all data, and one (red) excluding trees that died between 2017 and 2018.

**Fig. S4:**
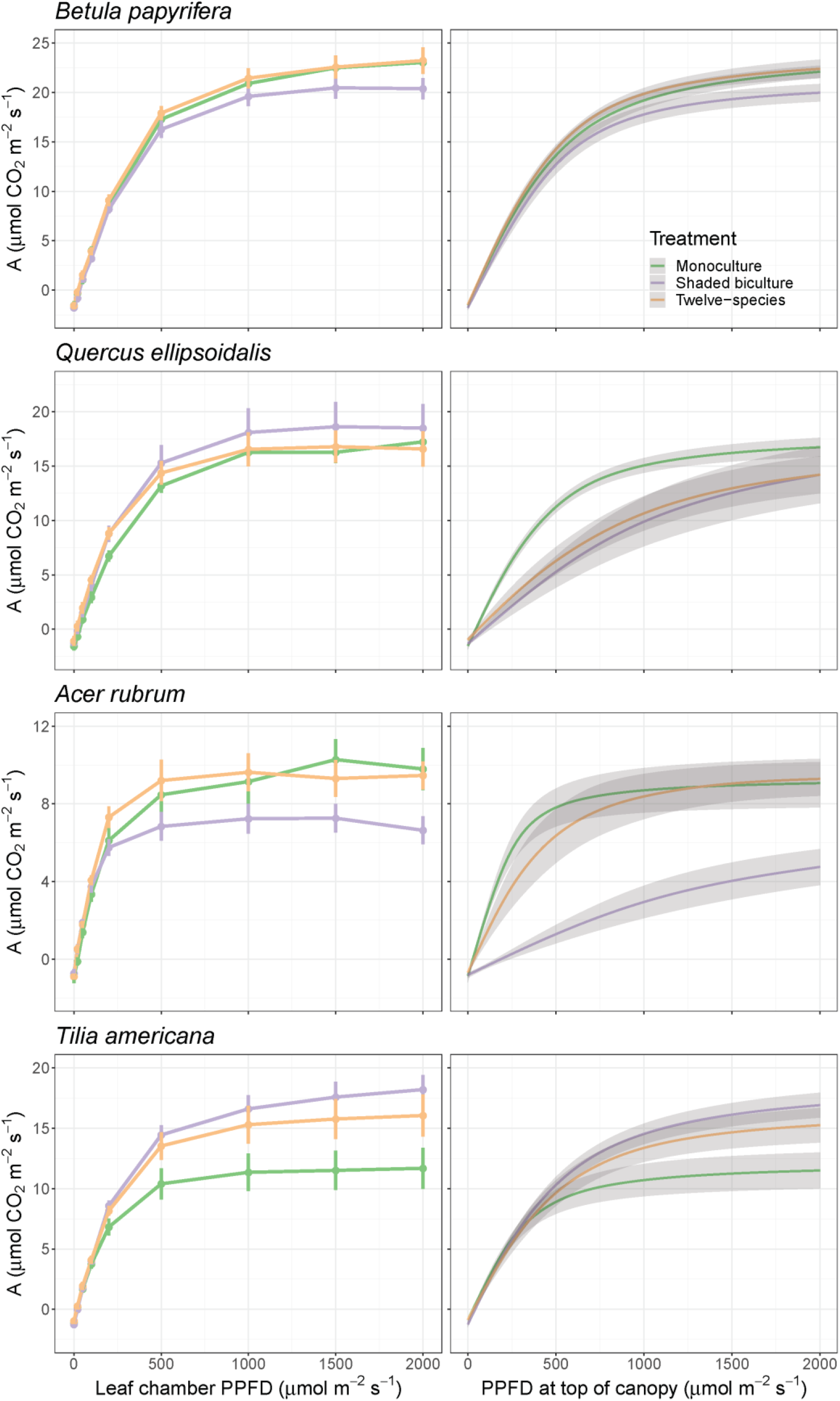
Area-based photosynthetic light-response curves for four broadleaf species. The left panels display carbon assimilation rates as a function of chamber light intensity; the right panels represent an estimate of realized assimilation rates as a function of light intensity at the top of the canopy. Error bars in left panels and gray ribbons in right panels are ± 1 SE.

**Fig. S5:**
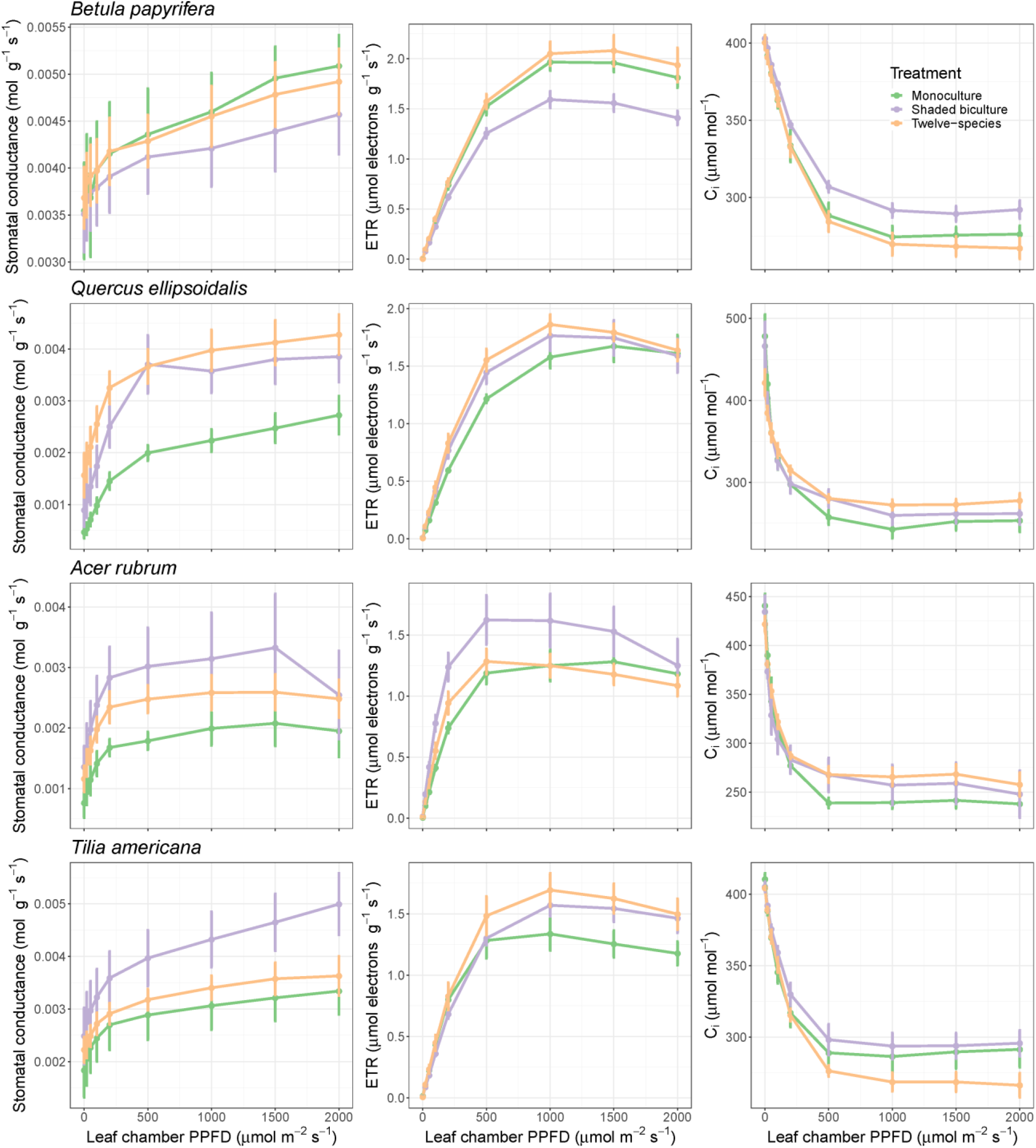
Responses of ETR, stomatal conductance, and *C*_*i*_ (internal CO_2_ concentration) to light availability for the four focal broadleaf species. Error bars are ± 1 SE.

**Fig. S6:**
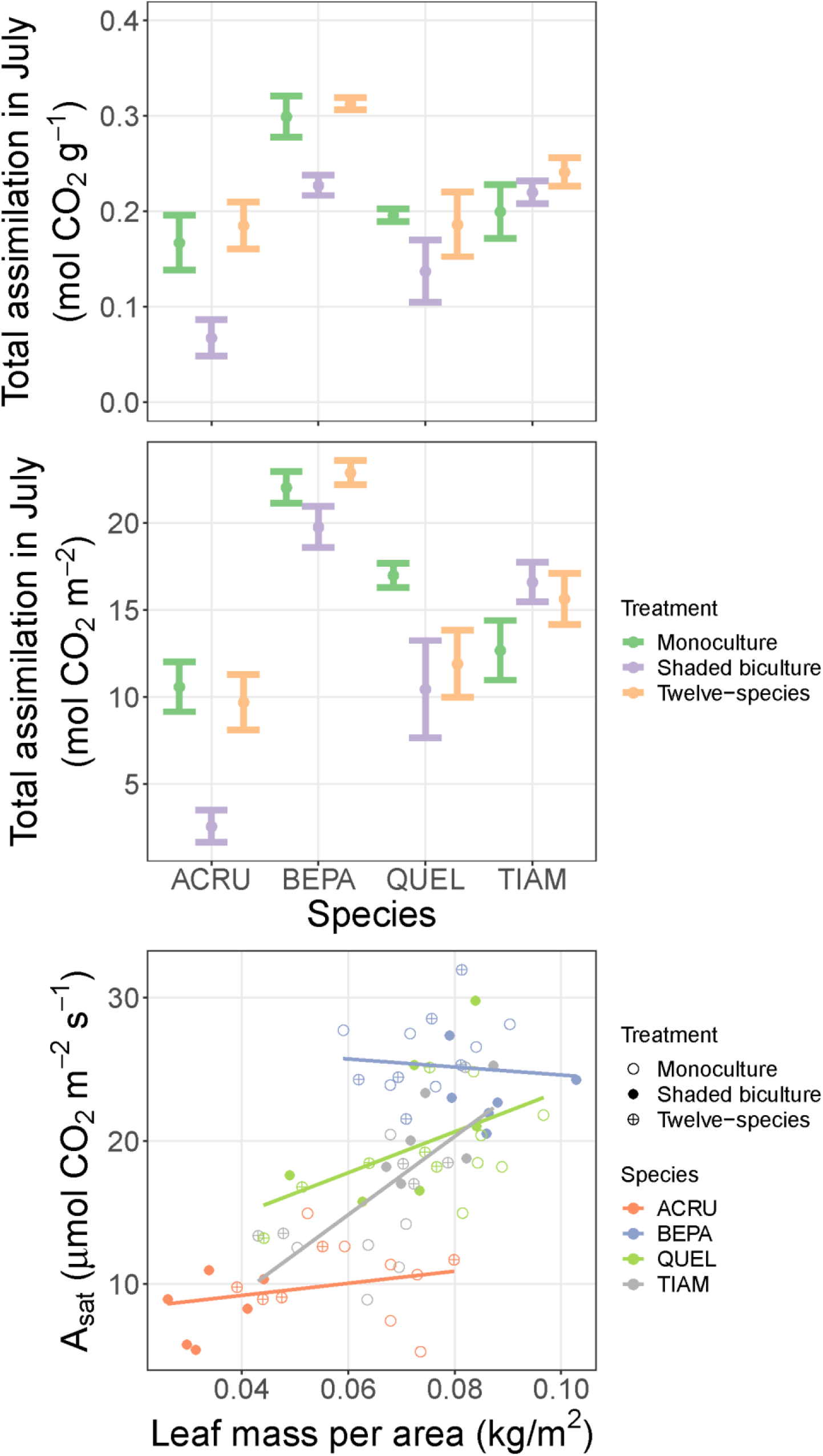
Estimates of total mass-based (top) and area-based (middle) carbon assimilation across species and treatments throughout the month of July, calculated by applying a time series of solar radiation from a nearby weather station as inputs to our rescaled light-response curves. Error bars are ± 1 SE. (bottom) Area-based *A*_*sat*_ is correlated with LMA across species; within species, there is only a positive relationship for *Quercus ellipsoidalis* and *Tilia americana*.

**Fig. S7:**
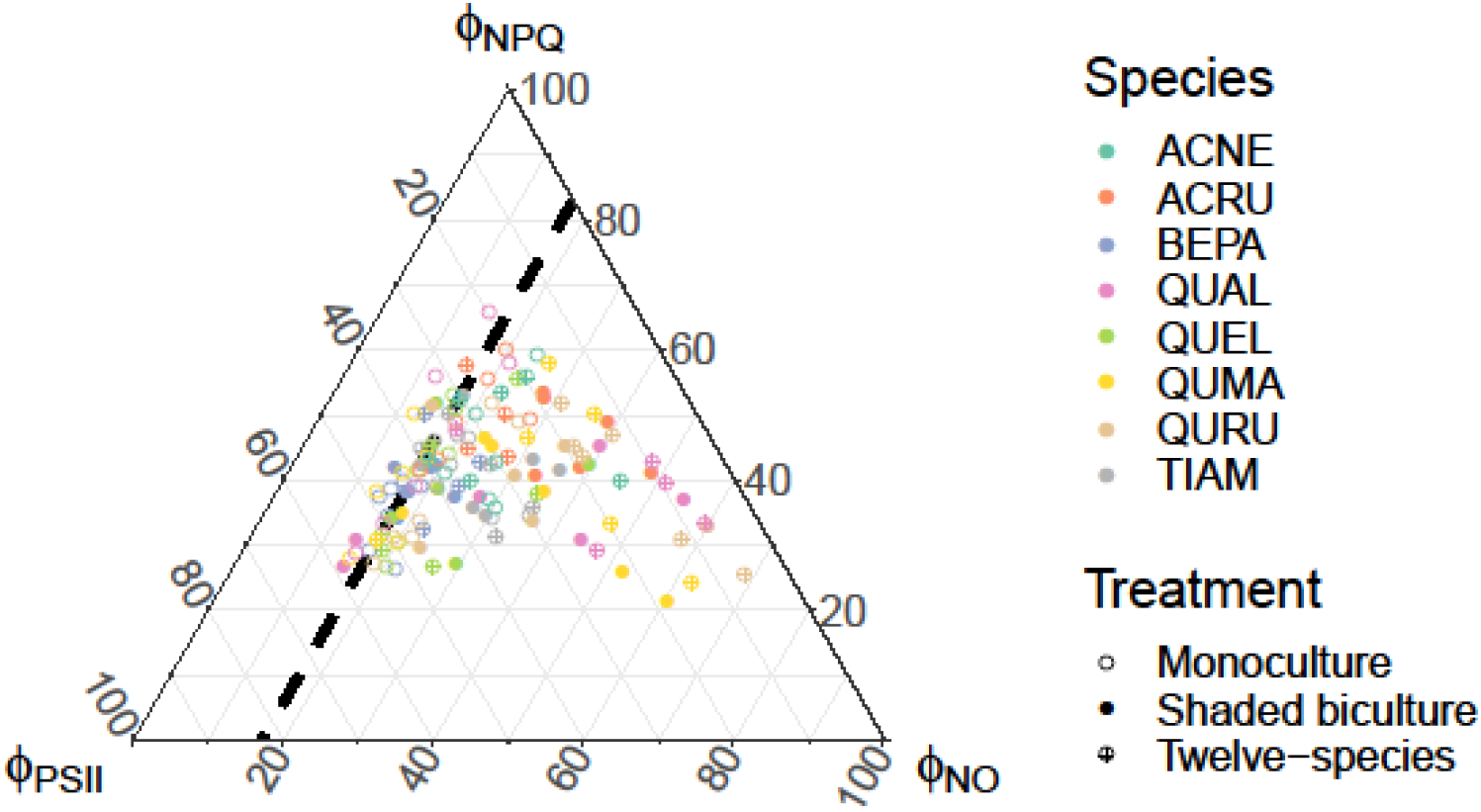
A ternary plot depicting variation among species and treatments in ϕ_PSII_, ϕ_NPQ_, and ϕ_NO_, as calculated using the lake model derivations of Kramer et al. (2004). These three quantities describe light energy dissipation through photochemistry, through non-photochemical quenching, and through non-regulated dissipation, respectively. A thick dashed line represents 0.83, the theoretical maximum efficiency of PSII posited in Tietz et al. (2017). Deviations to the lower right of this line represent increasingly high ϕ_NO_, which can result in photodamage.

**Fig. S8:**
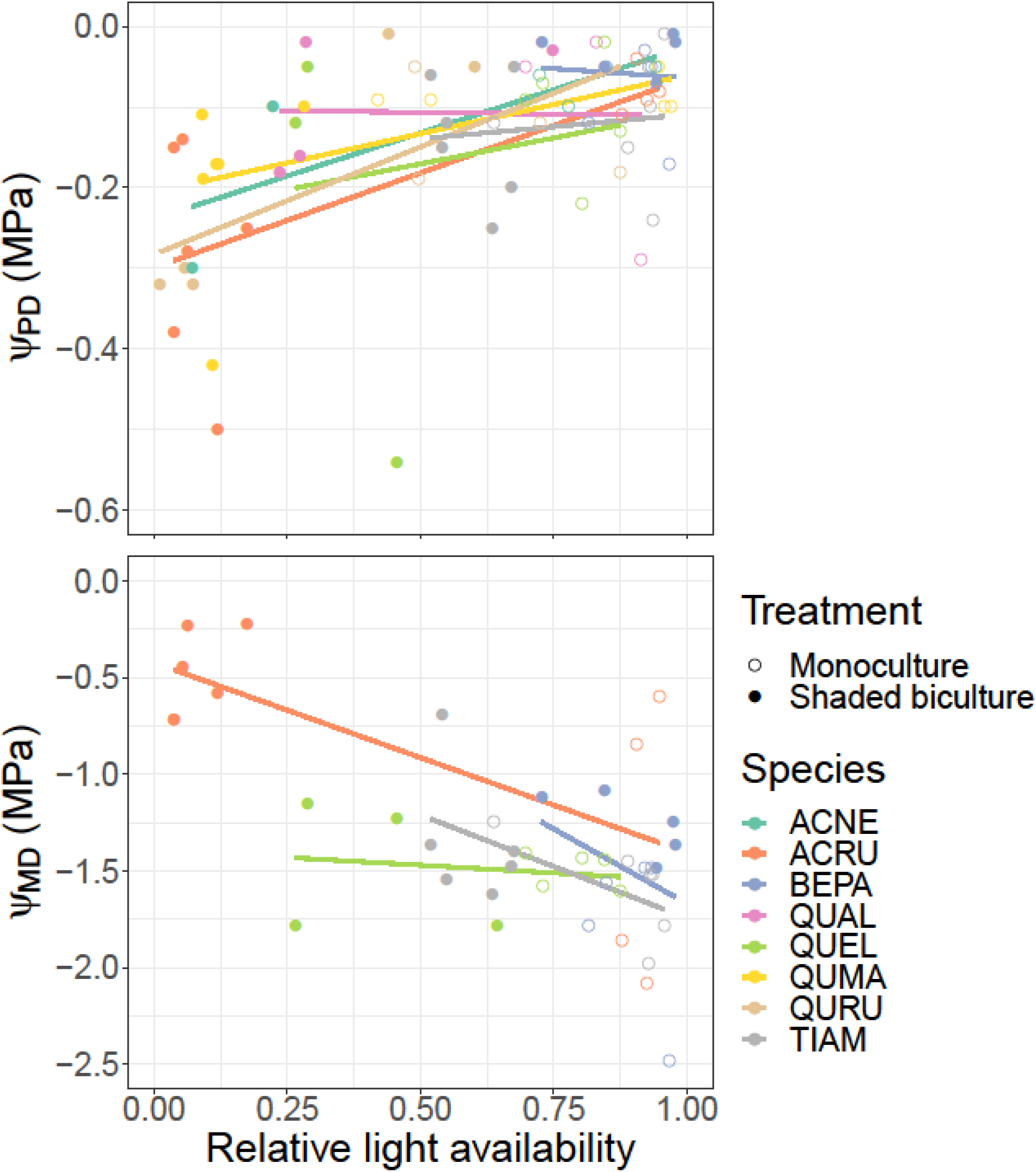
Water potential before dawn (Ψ_PD_; top) and at midday (Ψ_MD_; bottom) as a function of light availability. Data are from monoculture and shaded biculture treatments in all eight species (Ψ_PD_) or only the four focal species (Ψ_MD_). Best-fit lines come from species-specific OLS regressions. Two outliers in Ψ_PD_ are not included.

